# Transposable elements contribute to genome dynamics and gene expression variation in the fungal plant pathogen *Verticillium dahliae*

**DOI:** 10.1101/2021.01.25.428111

**Authors:** David E Torres, Bart PHJ Thomma, Michael F Seidl

## Abstract

Transposable elements (TEs) are a major source of genetic and regulatory variation in their host genome and are consequently thought to play important roles in evolution. Many fungal and oomycete plant pathogens have evolved dynamic and TE-rich genomic regions containing genes that are implicated in host colonization. TEs embedded in these regions have typically been thought to accelerate the evolution of these genomic compartments, but little is known about their dynamics in strains that harbor them. Here, we used whole-genome sequencing data of 42 strains of the fungal plant pathogen *Verticillium dahliae* to systematically identify polymorphic TEs that may be implicated in genomic as well as in gene expression variation. We identified 2,523 TE polymorphisms and characterize a subset of 8% of the TEs as dynamic elements that are evolutionary younger, less methylated, and more highly expressed when compared with the remaining 92% of the TE complement. As expected, the dynamic TEs are enriched in the dynamic genomic regions. Besides, we observed an association of dynamic TEs with pathogenicity-related genes that localize nearby and that display high expression levels. Collectively, our analyses demonstrate that TE dynamics in *V. dahliae* contributes to genomic variation, correlates with expression of pathogenicity-related genes, and potentially impacts the evolution of dynamic genomic regions.

**Significance statement:** Transposable elements (TEs) are ubiquitous components of genomes and are major sources of genetic and regulatory variation. Many plant pathogens have evolved TE-rich genomic regions containing genes with roles in host colonization, and TEs are thought to contribute to accelerated evolution of these dynamic regions. We analyzed the fungal plant pathogen *Verticillium dahliae* to identify TE variation between strains and to demonstrate that polymorphic TEs have specific characteristic that separates them from the majority of TEs. Polymorphic TEs are enriched in dynamic genomic regions and are associated with structural variants and highly expressed pathogenicity-related genes. Collectively, our results provide evidence for the hypothesis that dynamic TEs contribute to increased genomic diversity, functional variation, and the evolution of dynamic genomic regions.

## Introduction

Repetitive sequences such as transposable elements (TEs) are ubiquitous components of prokaryote and eukaryote genomes (Bowen and Jordan 2002; Wicker, et al. 2007). In eukaryotes, the repeat content can differ between species and range from only ∼5.6% in the fish *Tetraodon nigroviridis* (Jaillon, et al. 2004) to up to 85% in the maize genome (Schnable, et al. 2009; Jiao, et al. 2017). TEs can transpose to different locations within the host genome, and thus are often considered ‘mobile genetic elements’ (McClintock 1950). TEs have been traditionally classified into two classes based on their mode of transposition: class I retrotransposons transpose via a copy-and-paste mechanism, while class II DNA transposons mainly mobilize via a cut-and-paste system (Wicker, et al. 2007; Piégu, et al. 2015). TEs can impact host genome organization and gene function by disrupting protein-coding regions, modifying gene expression, or causing large-scale chromosomal rearrangements (Eichler and Sankoff 2003; Feschotte 2008; Bennetzen and Wang 2014; Le, et al. 2015; Mita and Boeke 2016). Consequently, TEs and their activities have been considered to generally decrease fitness, and TEs and TE-induced mutations are typically rapidly purged from a population. Furthermore, TE activities are generally suppressed by host defense processes. These processes include DNA methylation (Selker, et al. 2003; Zemach, et al. 2010), histone modifications associated with highly condensed heterochromatin (Slotkin and Martienssen 2007; Hocher, et al. 2018; Hocher and Taddei 2020), targeted processes such as RNA-silencing (Lippman and Martienssen 2004; Nicolas, et al. 2013; Yadav, et al. 2018), or repeat-induced point (RIP) mutation (Cambareri, et al. 1989; Galagan and Selker 2004; Fudal, et al. 2009; Wang, et al. 2020).

Over the last decades, besides their detrimental effects, TEs have been increasingly recognized to play important roles in adaptive genome evolution (Venner, et al. 2009; Hua-Van, et al. 2011; Lynch, et al. 2011; Bourque, et al. 2018). For example, TEs can be co-opted as a source of *cis*-regulatory elements (Chuong, et al. 2016; Kitano, et al. 2018) and mediate swift changes in the transcriptional landscape under biotic and abiotic stresses (Sinzelle, et al. 2009; Bourque, et al. 2018; Muszewska, et al. 2019). A classic example is the dark pigmentation of the peppered moth in response to industrialization that was caused by a single TE insertion into the first intron of a cell-cycle regulator gene, changing the transcript abundance of this gene and allowing the insect to camouflage on polluted surfaces (Cook and Saccheri 2013; Hof, et al. 2016). Changes in mutation rates in genes in proximity TEs can be caused by leakage of TE silencing mechanisms (e.g. RIP) or DNA repair following TE insertion (Fudal, et al. 2009; Rouxel, et al. 2011; Yoshida, et al. 2016; Wang, et al. 2020). Furthermore, abundant TEs and other repetitive elements can act as an ectopic substrate for double-strand break repair pathways, thereby fostering chromosomal rearrangements (structural variants; SVs), such as chromosomal translocations and inversions as well as large-scale duplications and deletions (Chan and Kolodner 2011; Faino, et al. 2016; Bourque, et al. 2018). For example, TE expansions in crops such as tomato, rice, wheat and soybean, have been shown to foster an increased accumulation of SVs associated with important traits during domestication (Fuentes, et al. 2019; Alonge, et al. 2020; Liu, et al. 2020; Walkowiak, et al. 2020). Interestingly, as TEs and SVs co-localize, they tend to form clusters in discrete regions in many eukaryotic organisms (Hastings, et al. 2009; Lin and Gokcumen 2019; Marand, et al. 2019). This co-localization, and its consequences for genome variation and organization, are important to understand the genome evolution of eukaryotic organisms (Kidwell and Lisch 2001; Wright and Finnegan 2001; Lynch and Conery 2003; Hurst, et al. 2004; Lynch 2007). Remarkably, these TE-rich regions are often enriched in environmentally responsive genes, for example in genes associated with roles in immunity or in secondary metabolism in plants (Sakamoto, et al. 2004; Kawakatsu, et al. 2016; Seidl, et al. 2016; van Wersch and Li 2019). Therefore, TE-rich compartments are considered relevant for contribution to adaptation to various abiotic and biotic stresses (Crombach and Hogeweg 2007; Knibbe, et al. 2007).

TE-rich compartments play important roles in the co-evolutionary ‘arms-race’ between pathogens and their hosts. Pathogens secrete molecules known as effectors to establish a successful parasitic relationship on their hosts (Rovenich, et al. 2014; Sánchez-Vallet, et al. 2018). In turn, hosts evolve receptors to activate immunity and halt pathogen colonization (Cook, et al. 2015; Han 2019). Thus, in order to overcome recognition by host immune systems, pathogens have to purge or diversify effector genes once their gene products become recognized (Moller and Stukenbrock 2017; Fouche, et al. 2018; Badet and Croll 2020). In many filamentous pathogens, effector genes are enriched in TE-rich and gene-sparse compartments, while they are typically underrepresented in TE-poor and gene-dense genomic regions that typically harbor housekeeping genes (Seidl and Thomma 2014; Dong, et al. 2015; Frantzeskakis, et al. 2020). TE-rich compartments are often characterized by increased substitution rates and increased occurrence of SVs and presence/absence polymorphisms (Raffaele, et al. 2010; Croll and McDonald 2012; de Jonge, et al. 2012; de Jonge, et al. 2013; Dutheil, et al. 2016; Hartmann and Croll 2017; Hartmann, et al. 2017; Wang, et al. 2017; Fokkens, et al. 2018; Plissonneau, et al. 2018; Grandaubert, et al. 2019; Gan, et al. 2020; Wyka, et al. 2020). Notably, a similar association of TEs with genes involved in immune responses has also been observed in plant hosts (Leister 2004; Kawakatsu, et al. 2016; Mascher, et al. 2017; Seidl and Thomma 2017). Therefore, TE-rich compartments have been suggested to facilitate the generation of variation that is necessary to fuel the ‘arms-race’ between pathogens and their hosts (Seidl and Thomma 2017; Frantzeskakis, et al. 2019; Schrader and Schmitz 2019; Frantzeskakis, et al. 2020; Torres, et al. 2020).

*Verticillium dahliae* is an asexual soil-borne fungal plant pathogen that colonizes the xylem of hundreds of susceptible host plants (Fradin and Thomma 2006; Klosterman, et al. 2009; Klosterman, et al. 2011; Inderbitzin and Subbarao 2014). Genomic comparisons between *V. dahliae* strains revealed that their genomes evolved by extensive chromosomal rearrangements (de Jonge, et al. 2013; Faino, et al. 2016). These rearrangements include chromosomal translocations, inversions, large-scale deletions, and duplications that collectively contributed to the formation of dynamic genomic compartments that originally have been referred to as lineage-specific (LS) regions, as these are variable between *V. dahliae* strains (Klosterman, et al. 2011; de Jonge, et al. 2012; de Jonge, et al. 2013; Faino, et al. 2016; Depotter, et al. 2019). Initially, LS regions were solely defined by presence/absence polymorphisms among sequenced strains (de Jonge, et al. 2013; Faino, et al. 2016). However, subsequent work on the chromatin landscape in *V. dahliae* has revealed that these LS regions display a unique chromatin profile (Cook, et al. 2020). Moreover, it was shown that a substantial number of additional genomic regions share chromatin characteristics with LS regions and, accordingly, that the amount of LS DNA has been underestimated (Cook, et al. 2020). The originally identified LS regions, together with the additional genomic regions that share a similar chromatin profile, are now referred to as dynamic genomic regions (Cook, et al. 2020). Importantly, these dynamic genomic regions are enriched for *in planta* induced effector genes that contribute to host colonization (Klosterman, et al. 2011; de Jonge, et al. 2012; de Jonge, et al. 2013; Kombrink, et al. 2017; Gibriel, et al. 2019; Cook, et al. 2020; Chavarro-Carrero, et al. 2021). Furthermore, these regions are enriched in relatively young and transcriptionally active TEs (Faino, et al. 2016; Cook, et al. 2020). While a key role of TEs to drive formation and maintenance of dynamic genomic regions has been revealed (Amyotte, et al. 2012; Faino, et al. 2015; Faino, et al. 2016; Shi-Kunne, et al. 2018; Depotter, et al. 2019; Cook, et al. 2020; Seidl, et al. 2020), it remains unclear what the impact of TE variation is on the biology of individual strains of *V. dahliae*. Changes in the TE landscape between *V. dahliae* strains, involving novel insertions as well as deletions, can cause structural as well as gene expression variation. Here, we exploited a collection of *V. dahliae* strains to analyze their TE landscape and systematically identify TE polymorphisms. Furthermore, we analyzed the polymorphic TE subset in detail to study its impact on genome function and organization. Our analysis extends previous hypotheses on the relevance of TE dynamics in dynamic genomic regions (Faino, et al. 2016; Cook, et al. 2020), and further characterizes a TE subset implicated in genomic variation and gene expression dynamics in *V. dahliae*.

## Results

### Structural variants are common in *V. dahliae*

Previous genomic comparisons among a narrow selection of *V. dahliae* strains have revealed extensive chromosomal rearrangements, which were proposed to have contributed to the establishment of dynamic genomic regions (de Jonge, et al. 2013; Faino, et al. 2016). To systematically assess their prevalence and abundance in a considerably larger collection of *V. dahliae* strains, we used paired-end sequencing reads for *de novo* prediction of SVs in 42 *V. dahliae* strains (Cook, et al. 2020; Chavarro-Carrero, et al. 2021) together with the gapless genome assembly of strain JR2 as a reference (Faino, et al. 2015). With a combination of four commonly used SV-callers (Supplementary Fig.1), 919 high-confidence SVs were identified in total, comprising 410 deletions, 49 inversions, 24 duplications, and 436 translocations (Figure 1A, Supplementary Fig.1). Importantly, we successfully recalled all 19 chromosomal rearrangements that were previously manually identified between *V. dahliae* strains JR2 and VdLs17 (Faino, et al. 2016). To investigate the occurrence of the 919 SVs across the *V. dahliae* strains, we calculated the frequency of each SV over all the strains included in the analysis. We observed that 87% (*n*=803) of the SVs is shared by <20% of the strains, i.e. they occur in less than eight strains (Figure 1B). Among the SVs, deletions occur at the highest frequency (Figure 1A, B; P<2×10^−16^, Kruskal-Wallis test). Interestingly, the number of SVs per strain does not correspond with the phylogenetic relationship among the *V. dahliae* strains (Figure 1C). Collectively, our data confirm that SVs occur commonly in *V. dahliae* and are typically shared by subsets of strains.

**Fig 1.**
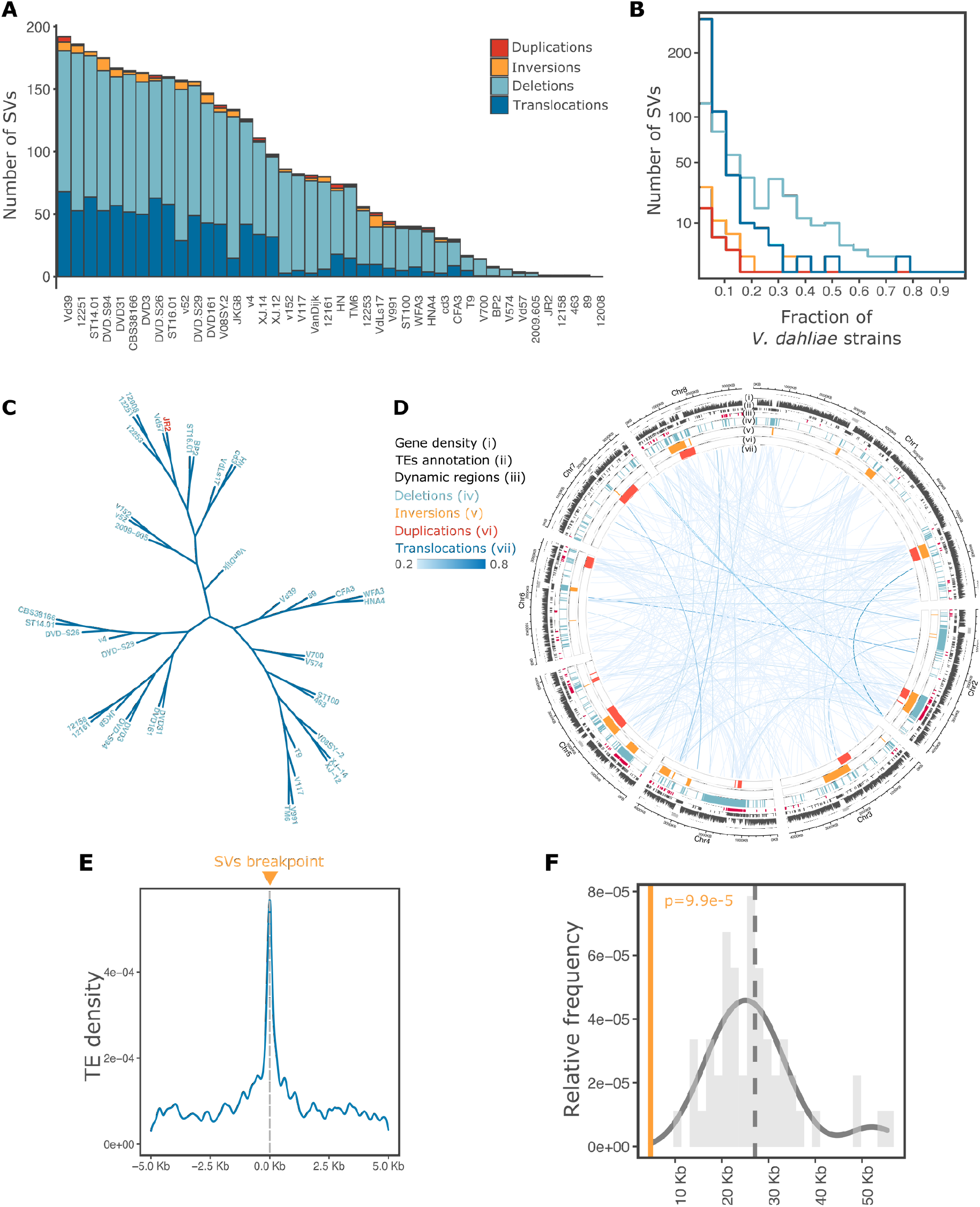
Structural variants (SVs) in a collection of 42 *Verticillium dahliae* strains. **(A)** The number of SVs is depicted per type for each *V. dahliae* strain included in the analysis; **(B)** Frequency distribution of the 919 SVs in the collection of *V. dahliae* strains, color-coded for variant type as indicated in panel **(A)**; **(C)** Unrooted neighbor-joining tree of the collection of *V. dahliae* strains based on 287,251 high-confident SNVs relative to the reference genome assembly of *V. dahliae* strain JR2 (red); **(D)** Circular plot displaying the genomic distribution of the 919 SVs along the eight chromosomes of the *V. dahliae* strain JR2 reference genome assembly. The tracks are shown in the following order from outside to inside: centromeric regions (Seidl, et al. 2020), gene density in 10 kb windows, TE distribution, dynamic genomic regions distribution (Cook, et al. 2020), 410 deletions (light blue), 49 inversions (yellow), 24 duplications (red), and 436 translocations (blue lines). The color intensity of the lines for an individual translocation is based on its frequency in the *V. dahliae* collection; **(E)** TE density around SV breakpoints is displayed in 10 kb windows; **(F)** The distribution of 10,000 random permutations of distances between SVs and TEs in dynamic genomic regions. The vertical dashed line represents the mean number of random occurrences, the yellow line indicates the average distance at 4,681.85 bp, significance P= 9.99×10^− 5^, z-score=−2.32.

Previously, we have demonstrated that SVs cluster at dynamic genomic regions in *V. dahliae* (de Jonge, et al. 2013; Faino, et al. 2016). Here, we similarly observed distinct genomic regions comprising multiple non-randomly distributed SVs (P<2×10^−16^, Kolmogorov-Smirnov test, Supplementary Fig.2), enriched in 17 regions (Pignatelli, et al. 2009) (hypergeometric test; P<0.05 after Benjamini-Hochberg correction; Supplementary Table 2), and co-localized with dynamic genomic regions (one-sided Fisher’s exact test, P = 0.00049; Supplementary Fig. 3). Interestingly, SVs occur largely independently of nucleotide mutations in dynamic genomic regions (Supplementary Table 3), confirming that SVs are a more likely phenomenon to increase variation in dynamic genomic regions than nucleotide mutations, especially when considering the previously reported high levels of sequence conservation in dynamic genomic regions (de Jonge, et al. 2013; Faino, et al. 2016; Shi-Kunne, et al. 2018; Depotter, et al. 2019).

### A subset of transposable elements in *V. dahliae* is polymorphic

TEs and other repetitive elements are often considered to drive the formation of SVs (Bourque, et al. 2018). In *V. dahliae*, TEs have been proposed to mediate genomic rearrangements (Faino, et al. 2016), and TEs and repetitive elements encompass 12.3% of the *V. dahliae* strain JR2 genome (Faino, et al. 2015). Class I LTR retroelements are the most abundant, while Class II DNA is less abundant (Supplementary Table 4). In line with previous observations (Klosterman, et al. 2011; de Jonge, et al. 2013; Faino, et al. 2015; Faino, et al. 2016; Cook, et al. 2020), TEs are significantly enriched (3.5-fold) in dynamic genomic regions when compared with the core genome and co-localize with SVs (Figure 1E-F; Supplementary Fig. 4). As TE dynamics may promote the formation of SVs, insertions, and deletions of TEs (i.e. polymorphic TEs) may directly impact host genome organization. To identify TE insertion and deletion polymorphisms, we used the paired-end sequencing data in the collection of 42 *V. dahliae* strains relative to the *V. dahliae* strain JR2 reference genome using TEPID (Stuart, et al. 2016). In total, we identified 2,387 deletions and 136 insertions, of which only 135 and 30 are unique deletion and insertion loci, respectively (Figure 2B). In the case of insertions, we decided to keep only elements that belong to designated TE superfamilies, removing twelve ‘unspecified’ elements. Only around 8% of all TEs are polymorphic, i.e. they show insertion or deletion events in different *V. dahliae* strains (Figure 2A). To further analyze the polymorphic TEs, we first calculated their frequency in the *V. dahliae* strains relative to strain JR2. This revealed that TE deletions occurred more frequently than insertions (P = 0.0349, one-sided Fisher’s exact test; Figure 2C). Only five TEs display both deletions and insertions, of which one belongs to the LTR/Gypsy and four to the LTR/Copia superfamily. Thus, only a limited subset of TEs in the *V. dahliae* genome is polymorphic, while the vast majority is shared by all strains analyzed.

**Fig 2.**
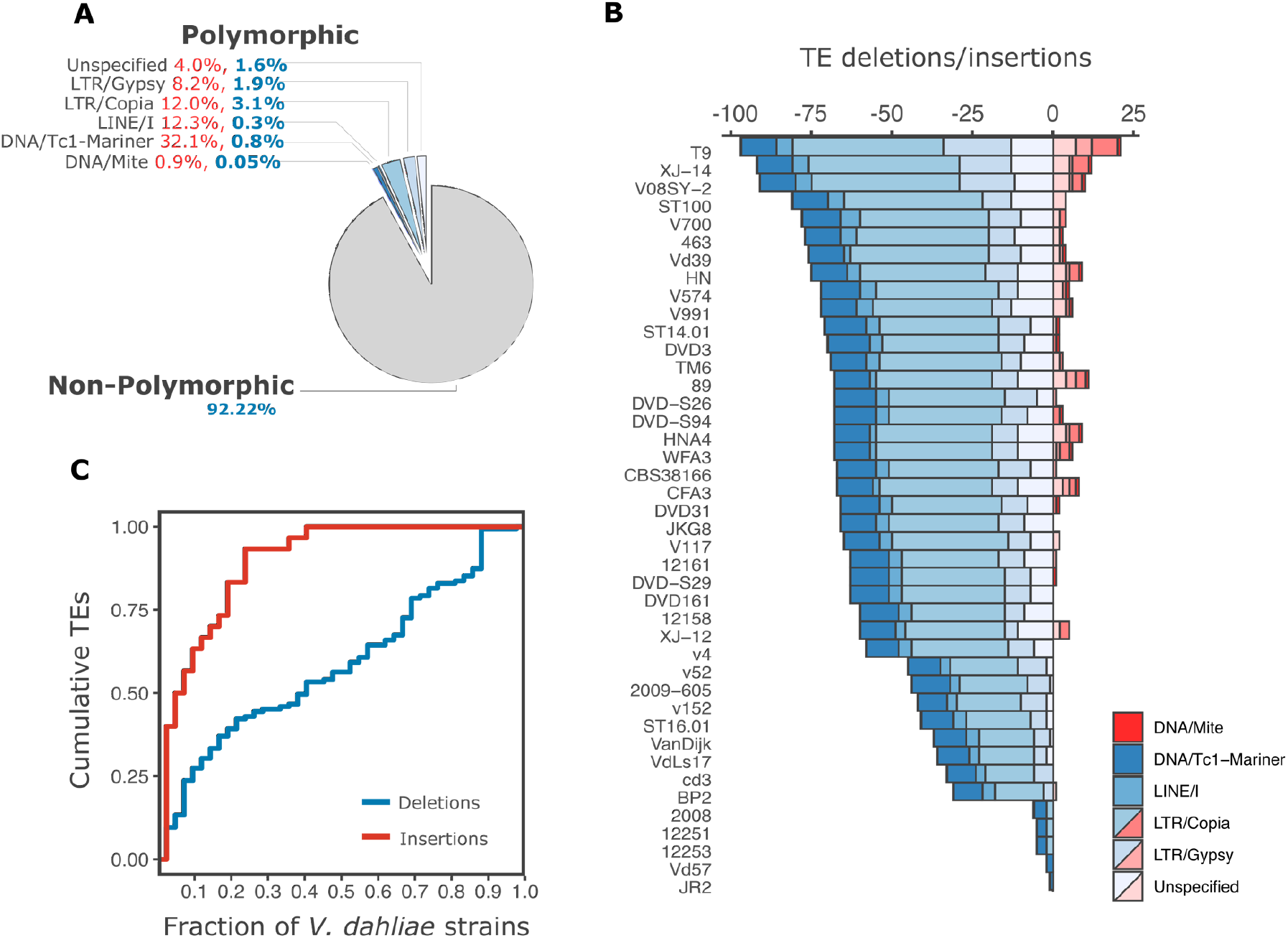
Transposable element polymorphisms in a collection of 42 *Verticillium dahliae* strains. **(A)** The percentage of polymorphic (shades of blue) and non-polymorphic (grey) TEs in *V. dahliae* strain JR2 is depicted as a pie chart. The percentages of polymorphic TEs separated by superfamily relative to the TE annotation of the *V. dahliae* strain JR2 is shown in blue, and the percentage of polymorphic TEs within each superfamily is shown in red; **(B)** The proportion of identified TE polymorphisms is shown for every *V. dahliae* strain, depicting deletions (blue) and insertions (red) per superfamily**; (C)** Cumulative frequency distribution of 165 polymorphic TEs in the *V. dahliae* strain collection, depicted by TE variant type.

### Polymorphic TEs are dynamic in *V. dahliae*

Fungi evolved a range of mechanisms to suppress TE activity and avoid TE expansions (Castanera, et al. 2016). However, we previously observed that a fraction of TEs is transcriptionally active in *V. dahliae*, suggesting it comprises TEs that are potentially able to multiply and accumulate in the *V. dahliae* genome (Faino, et al. 2016; Cook, et al. 2020). Therefore, we hypothesized that polymorphic TEs are not suppressed and still able to multiply. To assess if polymorphic TEs lack typical signatures of suppression when compared with non-polymorphic TEs in *V. dahliae* strain JR2, we assessed DNA methylation levels (in the three methylation contexts; mCG, mCHG, and mCHH), RIP signature (Composite Repeat-Induced-Mutations Index; CRI), Kimura distance (divergence from the TE consensus sequence), GC content, and expression levels *in vitro* (counts per million; CPM) by quantifying the abundance of TE-derived reads across all different TE copies in the genome using TEtranscripts (Jin, et al. 2015; Jin and Hammell 2018). Dimensional reduction by principal component analysis (PCA) revealed that polymorphic TEs cluster and are largely separated from the non-polymorphic TEs (Figure 3A). On the first dimension of the PCA, explaining 31.6% of the variation, polymorphic and non-polymorphic TEs are separated based on CRI, DNA methylation, and GC content (Supplementary Fig. 5). This observation can be explained by the association of TEs with strictly heterochromatic regions, in which TEs are characterized by high CRI and DNA methylation levels as well as by low GC content (Cook, et al. 2020). The second dimension of PCA, explaining 14.5% of the variation, revealed a more distinct separation between polymorphic and non-polymorphic TEs mainly based on Kimura distance, DNA methylation, and transcriptional activity *in vitro* (Supplementary Table 5). Specifically, we observed that the sequence of polymorphic TEs is more similar to the consensus sequence of the TE family (Kimura distance) and is characterized by a higher GC content when compared with non-polymorphic TEs, indicating that polymorphic TEs are evolutionary younger when compared with non-polymorphic TEs (Figure 3B). Notably, we also observed lower CRI and higher CG methylation levels, suggesting that polymorphic TEs are not silenced and are likely more active, as proxied by an increased transcriptional activity when compared with other TEs (Figure 3B). As expected, polymorphic TEs are expressed at higher levels *in vitro* and *in planta* when compared with non-polymorphic TEs (Figure 3B; Supplementary Fig. 6). Collectively, our analyses identified a subset of TEs as polymorphic among strains, and these tend to be evolutionary younger, less suppressed by RIP and methylation, as well as more highly expressed, and thus, we will now refer to polymorphic TEs as ‘dynamic TEs’.

**Fig 3.**
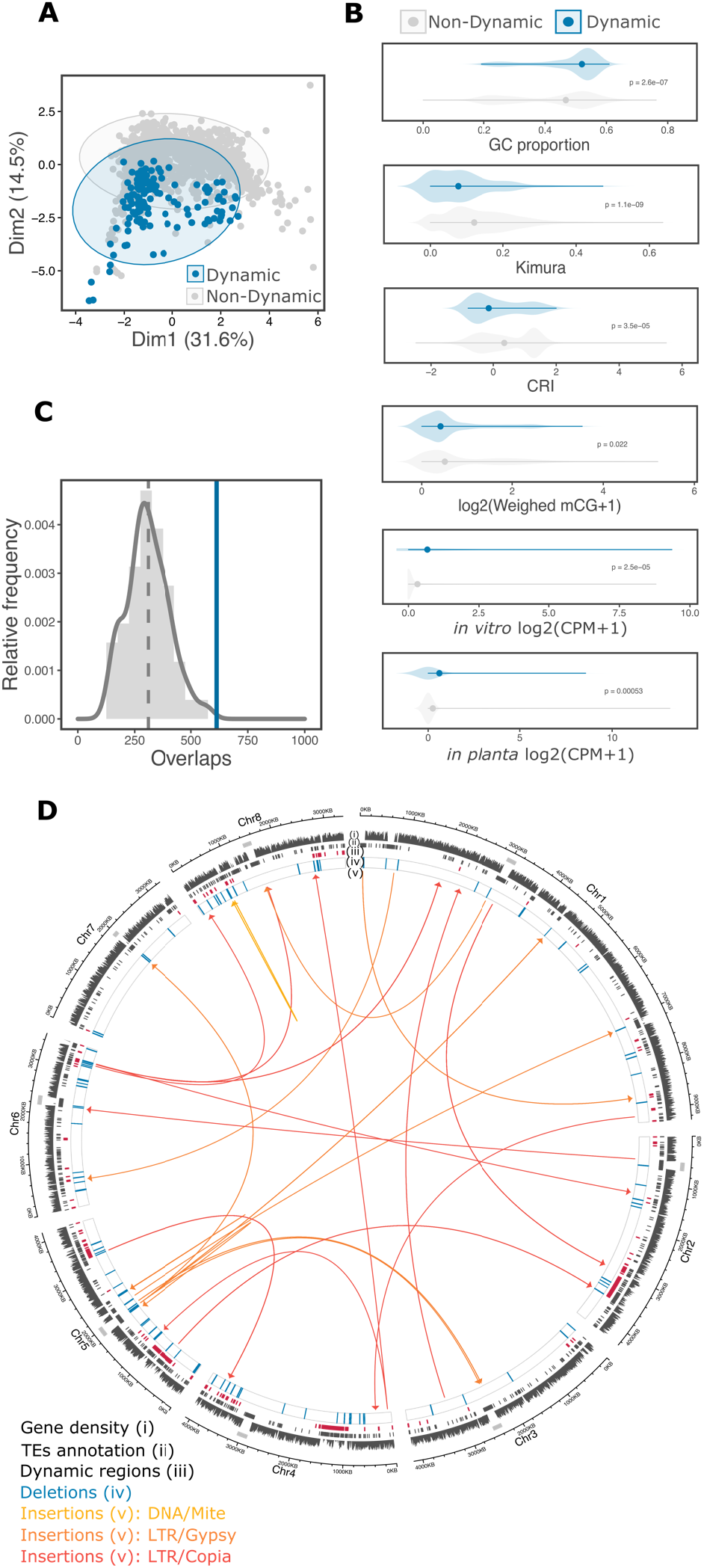
The TE landscape across the *V. dahliae* genome is dynamic. **(A)** Principal Component Analysis of TEs based on methylation (mCG, mCHG, mCHH), GC proportion, Kimura distance, Composite Repeat Induced Point Mutation index (CRI), frequency, and *in vitro* expression. Each point represents a single TE and colored ellipses represent the confidence interval for the dynamic and non-dynamic TEs; **(B)** Dynamic (*n*=165) and non-dynamic TEs (*n*=1,956) were compared based on GC proportion, the Kimura distance to the TE family consensus sequence, CRI, cytosine methylation, and *in vitro* expression (CPM; Counts per Million). Statistical significance was assessed using one-sided Wilcoxon rank-sum test; **(C)** The association between polymorphic TEs and dynamic genomic regions was tested using a permutation test. The vertical dashed line shows the mean number of polymorphic TEs overlaps expected at 10,000 random permutations and the blue line indicates the empirical number of overlaps, significance P= 9.99×10^−5^, z-score=3.2927; **(D)** Circular plot displays the genomic distribution of 165 dynamic TEs along the eight chromosomes of *V. dahliae* strain JR2. The tracks display the centromeric regions, gene density (in 10 kb windows), TE annotation, dynamic genomic regions, TE deletions, and TE insertions (with arrows indicating the insertion direction) from outside to inside.

Dynamic genomic regions in *V. dahliae* have been previously shown to be enriched for transcriptionally active TEs (Supplementary Fig. 4; (Faino, et al. 2016; Cook, et al. 2020)). We hypothesized dynamic genomic regions are enriched for dynamic TEs. Therefore, we analyzed the distribution of dynamic TEs in *V. dahliae* strain JR2 (Figure 3D), leading to the identification of clusters of dynamic TEs that co-localize with dynamic genomic regions (P=0.00024, one-sided Fisher’s exact test; Supplementary Table 6; Figure 3C). Interestingly, dynamic TEs do not localize closer to SVs than non-dynamic TEs, suggesting that both types of TEs can contribute to the formation of SVs. While deleted TEs were enriched in dynamic genomic regions, we could not observe a similar enrichment for TE insertions in the JR2 strain (Supplementary Fig. 7). Dynamic TEs belonging to LTR/Copia superfamilies or unspecified repetitive elements are highly enriched in dynamic regions (P=8.24×10^−9^ and P=1.36×10^−5^ respectively, one-sided Fisher’s exact test), and for instance, LTR/Copia represents 51.1% of all dynamic TEs in dynamic regions (Supplementary Table 7). Therefore, our results suggest that the LTR superfamily is an important component of the dynamic TE landscape.

### Dynamic TEs impact the functional genome of *V. dahliae*

Next to their association with SVs, TE presence and activity have been suggested to directly or indirectly impact the functional genome, for example by modifying the expression of genes in their vicinity or by inducing changes in gene structure (Bourque, et al. 2018; Schrader and Schmitz 2019). To establish to which extent TEs may affect gene expression in *V. dahliae*, we first summarized the occurrence of TEs up to 5 kb upstream and downstream of all protein-coding genes (Supplementary Fig. 8). The majority of annotated TEs is located within a 5 kb range of a gene (67% of total TEs, *n*=1,418; P=0.0094, one-sided Fisher’s exact test), with an overrepresentation of dynamic TEs belonging to the LTR/Copia, LINE/I, and DNA/Tc-1-Mariner superfamilies (Figure 4A; Supplementary Fig. 8). By assessing the expression of TEs as a proxy for their activity, we observed that many families are expressed *in vitro* and typically at higher levels during infection (P<0.05, Figure 4B; Supplementary Fig. 9). As expected, these families are abundant in dynamic TEs (Figure 4B). Thus, environmental changes potentially induce the expression of TEs.

**Fig 4.**
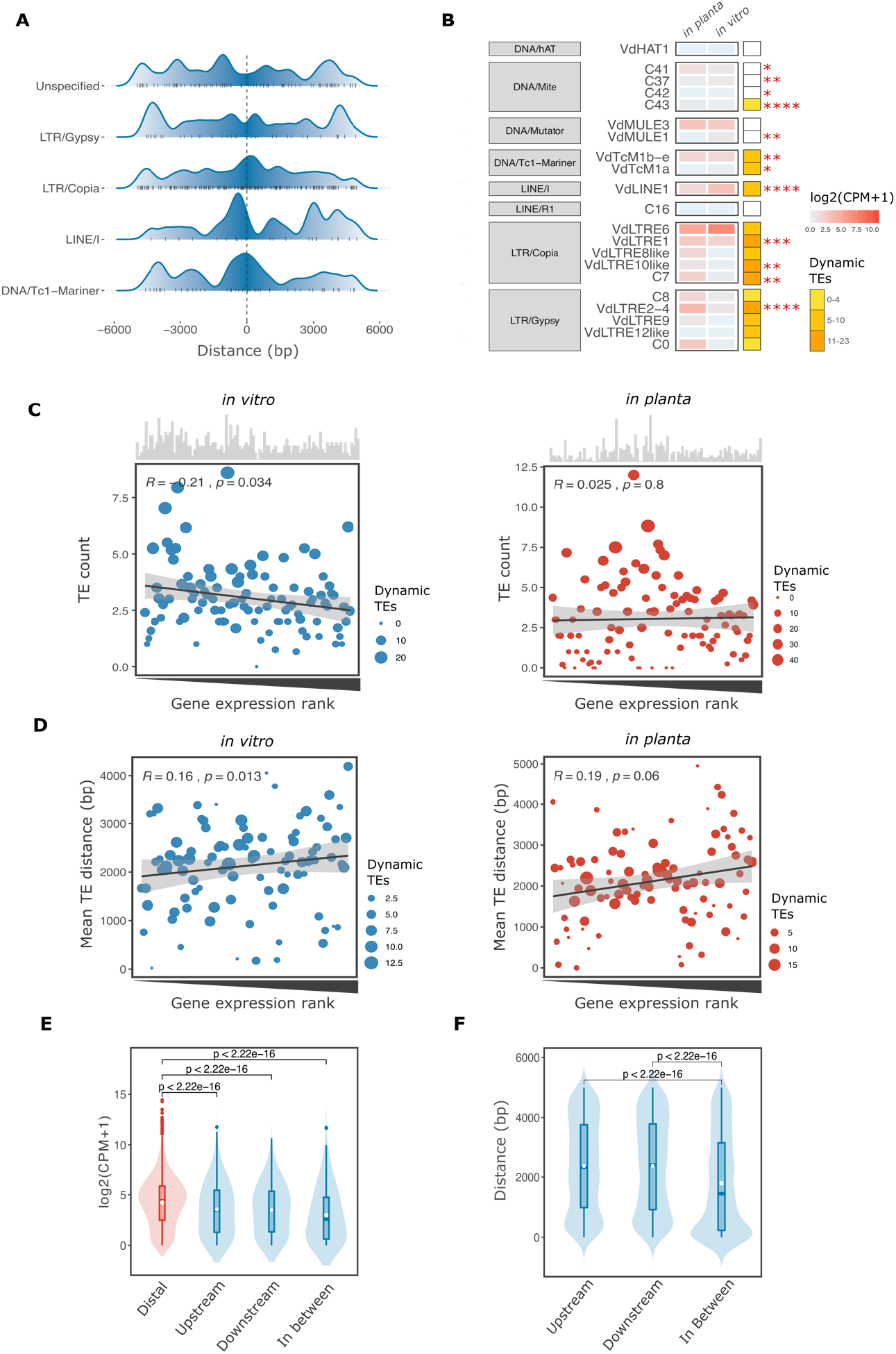
The presence of dynamic transposable elements is correlated with expression of genes that localize nearby. **(A)** Ridgeline shows the density distribution of dynamic TEs, classified by TE superfamily, upstream and downstream of genes; **(B)** Unsupervised clustering of TE expression per TE family and superfamily, color-coded based on the mean-per-family expression value (log2(CPM+1)). The column depicts the abundance of dynamic TEs within each group (yellow coded). Statistical significance for comparisons between *in vitro* (growth in MS media) and *in planta* (colonization of *Arabidopsis thaliana* at 28 days post inoculation) was assessed using a one-sided Wilcoxon rank-sum test * P<=0.05, ** P<=0.01, *** P<=0.001, **** P<=0.0001; **(C)** Relationship of gene expression ranks *in vitro* (blue) and *in planta* (red) and the number of TEs. Low rank = low expression and high rank = high expression. The plots display linear regression (dark line) and confidence interval (light grey) as well as the R and P values after linear regression. The histograms on top of the graph show the number of dynamic TEs per expression rank and the dot-size reflects the proportion of dynamic TEs relative to the total number of TEs per expression rank; **(D)** Relationship of gene expression rank and mean distance to TEs (bp) with annotations as for **(C)**; **(E)** Gene expression in relation to the TE context (TEs within 5 kb); no TE in proximity = Distal (*n*=8,742), or TEs flanked upstream (*n*=1,098), downstream (*n*=1,051), or in between (*n*=546; blue). Significant differences were assessed using one-sided Wilcoxon rank-sum test; **(F)** TE distance to genes in 5 kb windows; TEs in upstream (*n*=1,609), downstream (*n*=1,511) or in between (*n*=1,582). Statistical differences were assessed using one-sided Wilcoxon rank-sum test.

To further assess the correlation between the proximity of TEs and the expression of nearby genes, we compared the expression of genes located within 5 kb windows from TEs under *in vitro* and *in planta* conditions. We observed that 23% of all annotated genes reside within 5 kb of a TE (*n*=2,696). Genes in proximity to TEs are transcribed at significantly lower levels when compared with genes that do not reside in close proximity of TEs (P<2×10^−16^, *in vitro*, P=1.9×10^−9^ *in planta*; Kruskal-Wallis test). Therefore, we hypothesized that the density of dynamic TEs correlates with expression in *V. dahliae*. To investigate this, we aggregated genes and TEs within 5 kb by their expression rank and obtained the number of TEs in each rank. We observed only a moderate negative correlation between the number of dynamic TEs and gene expression ranks *in vitro* (Figure 4C; R = –0.21, P = 0.034 linear regression). Conversely, the distance between genes and dynamic TEs is moderately positively correlated with higher expression ranks (Figure 4D; R = 0.16, P = 0.013 linear regression). These results suggest that dynamic TEs are associated, albeit weakly, with genes expressed at low levels under *in vitro* conditions. However, we observed a different pattern when considering host colonization, as gene expression ranks *in planta* did not correlate with TE density (Figure 4C; R = 0.025, P = 0.8 linear regression) whereas, similar to *in vitro* conditions, TE distance and expression are moderately positively correlated (Figure 4D; R = 0.19, P = 0.06 linear regression). Collectively, our data suggest the dynamic TE density nearby protein-coding genes correlates only weakly with gene expression in different conditions.

To assess the relationship between TEs and gene expression, we categorized genes into four categories based on the relative position of TEs; genes with no TEs in proximity (*n*=8,742), genes with TEs localized upstream (*n*=1,098), downstream (*n*=1,051), or genes flanked by TEs on both sides (*n*=546). The distance between TEs and genes is significantly shorter when genes are flanked by TEs, when compared with genes having TEs only upstream or downstream (Figure 4F). Importantly, genes flanked by TEs show reduced expression *in vitro* when compared with genes in the proximity of TEs located either upstream or downstream (Figure 4E), a pattern we similarly observed during host colonization. We similarly observed this pattern when only considering the subset of dynamic TEs (Supplementary Fig. 10, Supplementary Fig. 11). Interestingly, during host colonization genes in proximity to dynamic TEs do not show a significant decrease in expression (Supplementary Fig. 10). Thus, genes in proximity to non-dynamic TEs showed lower expression levels when compared with genes in proximity to dynamic TEs.

### Pathogenicity-related genes co-localize with dynamic TEs

To understand which types of genes are enriched in proximity to TEs, we performed functional annotation of the *V. dahliae* strain JR2 gene catalog using a Cluster for Orthologs Groups (COG) approach and subsequently tested for enrichment of genes with specific COG categories. No significant enrichment of genes in proximity to TEs could be observed for most of the categories belonging to ‘cellular processes’ or to ‘information processing’. In contrast, genes annotated as ‘metabolism’ and ‘poorly characterized’ are significantly enriched (Figure 5A). Especially, genes belonging to the sub-categories ‘defense mechanisms’ or ‘carbohydrate metabolism’ are highly enriched (Figure 5A). A total of 39% of genes that reside in proximity to TEs fall in the category ‘function unknown’ (*n*=579) or ‘not categorized’ (*n*=482), which refers to orthologs present in other organisms but without known function. Furthermore, we could not find orthologs for 14.9% of the genes that reside in proximity to TEs (*n*=402), described in this work in the ‘not recognized orthologs’ category, which can probably be explained as gene content specific to the *V. dahliae* lineage (Figure 5A; Supplementary Table 8). Interestingly, most of these gene categories are associated with TEs (Figure 5A) and overlap with various SVs (Figure 5B). Subsequently, we annotated all protein-coding genes for functions that have previously been associated with pathogenicity, such as genes encoding secreted proteins (*n*=672), carbohydrate-active enzymes (CAZymes, *n*=498), secondary metabolites (*n*=25), or effector candidates (*n*=193). These genes can be broadly classified as ‘pathogenicity-related’ and are enriched in the ‘poorly characterized’ functional category, especially secreted proteins and effector candidates (P=3.49×10^−15^ and *n*=390, P=2.85×10^−6^ and *n*=127, respectively, one-sided Fisher’s exact test; Supplementary Table 9). Genome-wide, we observed that 40.72% (*n*=401) of predicted pathogenicity-related genes have a TE localized within 5 kb (Figure 5E). Dynamic TEs are enriched in proximity (5 kb) to effector candidates and secondary metabolites (P=0.0080 and P=0.0186, respectively; one-sided Fisher’s exact test), even though we did not observe that these genes are significantly closer to dynamic TEs (Figure 5G). Thus, commonly considered pathogenicity-associated genes reside in proximity to TEs, yet only some genes are in proximity to dynamic TEs.

**Fig 5.**
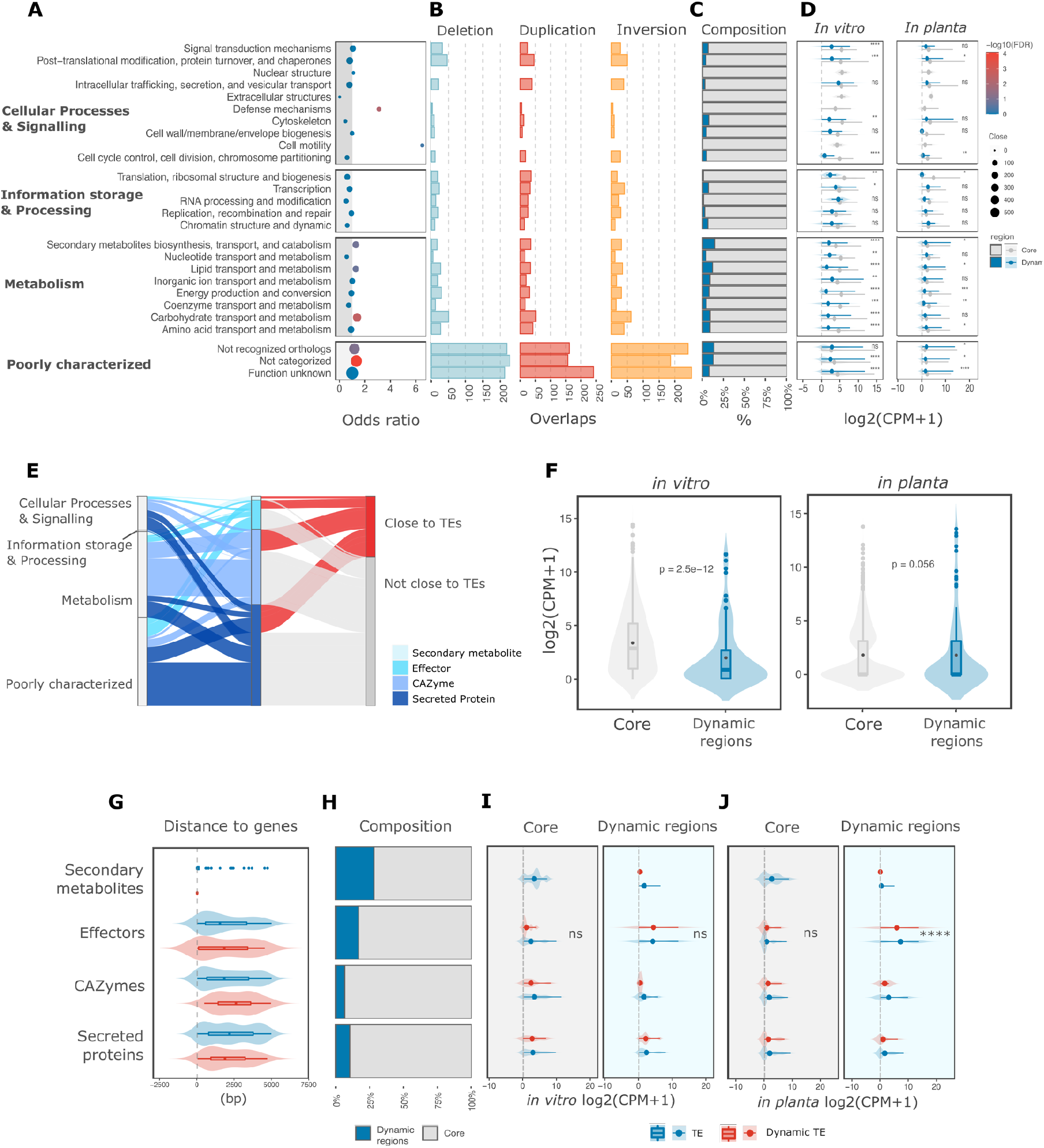
Dynamic transposable elements correlate with expression of a subset of genes. **(A)** Enrichment of TEs in proximity (within 5 kb range) to genes in different functional categories. Functional annotation was based on Clustering for Orthologous Groups (COGs) categories. The y-axis shows the different functional categories and the x-axis the odds ratio after one-sided Fisher’s exact test; the grey shadow represents values below 1 odds ratio, and the -log10(P-value) after Benjamini-Hochberg correction for false-discovery rate is color-coded. The size is relative to the number of genes associated with TEs in each category; **(B)** The number of genes overlapping deletions (blue) and duplications (red), and inversions (yellow); **(C)** The relative proportions of genes localized in the dynamic genomic regions (blue) and core (grey) genomic regions; **(D)** Gene expression distribution in two conditions (*in vitro* and *in planta*; log2(CPM+1)) and separated by core and dynamic genomic regions; **(E)** Predicted pathogenicity-related genes (*n*=1,388) overlapping with COG categories and depicting the proportion of genes with TEs in proximity (5 kb); **(F)** Gene expression of pathogenicity-related genes in two conditions (*in vitro* and *in planta*; log2(CPM+1)) and separated by core and dynamic genomic regions; **(G)** Distance between genes and TEs separated by dynamic and non-dynamic TEs. Statistical significance was assessed using Wilcoxon rank-sum test; **(H)** The composition of pathogenicity-related gene categories separated in dynamic (blue) and core (grey) genomic regions; Pathogenicity-related gene expression *in vitro* **(I)** and *in planta* **(J)** separated by dynamic and non-dynamic TEs. Statistical significance was assessed by a Kruskall-Wallis rank-sum test; ****P<=0.0001.

When compared with the core genome, dynamic genomic regions in *V. dahliae* are relatively gene-poor (Figure 5C) and SV-rich (Supplementary Fig. 3), as well as enriched for *in planta*-expressed genes (de Jonge, et al. 2012; de Jonge, et al. 2013; Kombrink, et al. 2017; Cook, et al. 2020; Chavarro-Carrero, et al. 2021) (Figure 5D). While a broad range of functional categories can be observed in dynamic genomic regions (Figure 5C), these are enriched for genes encoding secreted proteins and candidate effector genes (Figure 5H; (Cook, et al. 2020); P=0.032 and P=0.014, respectively; one-sided Fisher’s exact test). Since we have shown that gene expression levels correlate with the presence or depletion of TEs (Figure 4), we queried to which extent dynamic TEs in proximity of pathogenicity-related genes correlate with gene expression patterns in the dynamic genomic regions. We observed a significant enrichment of dynamic TEs co-localizing with highly expressed genes *in vitro* and *in planta* (P=0.0116 and P=0.0010 for 50% and 75% upper quartiles for expression, respectively, after one-sided Fisher’s exact test; Supplementary Table 10). Under *in vitro* conditions, pathogenicity-related genes located in the core genome are expressed significantly higher than genes located in dynamic genomic regions (Figure 5F). As expected, we observed the opposite trend *in planta* (Figure 5F), where pathogenicity-related genes that are localized in dynamic genomic regions are highly expressed during host colonization (de Jonge, et al. 2012; de Jonge, et al. 2013). Furthermore, we observed an enrichment of dynamic TEs located within 5 kb of genes of dynamic genomic regions (P=0.0062 after one-sided Fisher’s exact test). As expected, a higher proportion of these highly expressed genes are effector candidates (Figure 5I, J). These results suggest that dynamic TEs occur more likely in close proximity to highly expressed pathogenicity-related genes in dynamic genomic regions.

## Discussion

Transposable elements (TEs) can contribute to genomic and transcriptomic variation (Bourque et al., 2018) and, consequently, TEs have been often considered to play crucial roles in genome evolution and function (Feschotte 2008). Here, we analyzed the TE landscape in the plant pathogen *V. dahliae*. We show that the presence of TEs is associated with abundant SVs that emerged independently in individual *V. dahliae* strains. The SVs form discrete clusters and co-localize within dynamic genomic regions. Importantly, we demonstrate that these SVs co-locate with only a subset of TEs that can be defined as dynamic, considering that they are polymorphic, evolutionary younger, less silenced, and higher expressed. Furthermore, we show that the presence of dynamic TEs is associated with changes in gene expression of genes that can be assigned to diverse functional categories, particularly with host-pathogen interaction genes. Collectively, our results provide evidence for the hypothesis that dynamic TEs contribute to increased genomic diversity, functional variation, and the evolution of dynamic genomic regions.

TEs occur commonly in eukaryote genomes, yet there is an enormous variation in the TE content and diversity between different organisms (Wicker, et al. 2007; Huang, et al. 2012; Dietrich, et al. 2013). Typically, TE expansions are suppressed by various genome defense mechanisms, but some TE copies can escape suppression and remain active and mobile. Polymorphic TEs have been associated with recent transposition activity (Huang, et al. 2012), and in *V. dahliae*, a subset of TEs shares these characteristics. Only a subset of all predicted TEs is young and dynamic, which is in contrast to some other fungi in which there is a considerable amount of young TEs associated with recent TE expansions, such as in *Zymoseptoria tritici* (Badet, et al. 2020; Lorrain, et al. 2020; Oggenfuss, et al. 2020), *Leptosphaeria maculans* (Grandaubert, et al. 2014), *Blumeria graminis* (Frantzeskakis, et al. 2018) and *Magnaporthe oryzae* (Kang, et al. 2016). In many plants and fungi, TE expansions are mainly driven by Long Terminal Repeat (LTR) retrotransposons (Vitte and Bennetzen 2006; Tsukahara, et al. 2009; Muszewska, et al. 2011; Huang, et al. 2012; Amselem, et al. 2015; Donnart, et al. 2017; Galindo-Gonzalez, et al. 2017; Lorrain, et al. 2020). We observed an enrichment of LTRs in facultative heterochromatic regions and at the centromeres in *V. dahliae* (Cook, et al. 2020; Seidl, et al. 2020). Similarly, LTRs are an important fraction of the dynamic TEs in *V. dahliae* that are localized in the dynamic genomic regions. TE expansions have occurred at least twice during the evolution of *V. dahliae*, once during *Verticillium* diversification, and then right after *V. dahliae* speciation (Faino, et al. 2016). The active dynamic TEs we identified are likely the product of the most recent TE expansion in *V. dahliae*, and likely contributed to the formation of the dynamic genomic compartments.

Dynamic TEs are a small fraction of the total TE repertoire, which suggests that most TEs are suppressed. Three main processes have been associated with TE suppression in fungi: repeat-induced point mutation (RIP), DNA methylation, and RNA-mediated silencing. RIP is usually associated with sexual fungal organisms (Galagan and Selker 2004), yet RIP mutations have been also observed in *V. dahliae* that is considered asexual (Clutterbuck 2011; Klosterman, et al. 2011; Amyotte, et al. 2012; Cook, et al. 2020). RIP mutations have been associated with meiosis, but alternative mechanisms during vegetative propagation could lead to RIP-like mutations (Clutterbuck 2011). This has been recently observed in *Z. tritici* and *N. crassa* in which propagation through mitotic divisions generate RIP-like mutations (Möller, et al. 2020; Wang, et al. 2020). Intriguingly, dynamic TEs are depleted in RIP mutations. Consequently, we consider different RIP scenarios, such as that RIP activity was lost during an evolutionary transition to asexuality in *V. dahliae*. In such a scenario, the mutations we observed in most TEs are remains of ancestral RIP activity. A second scenario is that *V. dahliae* is not strictly asexual, but rare sexual activity occurs in the population. In this scenario, the RIP machinery may be active during the occasional meiosis. Thirdly, a RIP-like mechanism could act outside the typical meiotic cycles and might be active in *V. dahliae*. These hypotheses are not mutually exclusive, and it remains challenging to elucidate the mechanistic origin of RIP mutations in *V. dahliae*.

Next to RIP, DNA methylation targets repetitive elements in many fungi (Nakayashiki, et al. 1999; Zemach, et al. 2010; Montanini, et al. 2014; Bewick, et al. 2019). We have previously demonstrated that the majority of TEs in *V. dahliae* is methylated (Cook, et al. 2020; Seidl, et al. 2020). Intriguingly, we now show that dynamic TEs are less methylated and are do not show signs of RIP, collectively indicating that these elements are not suppressed. In the model plant *Arabidopsis* and the fruit-fly *Drosophila*, polymorphic and active TEs occur in discrete regions where novel insertions are not efficiently suppressed, while TE insertions outside of these regions are mostly suppressed by methylation (Hollister and Gaut 2009; Tsukahara, et al. 2009; Stuart, et al. 2016; Lee and Karpen 2017). We observed that TEs in dynamic genomic regions display low methylation levels (Cook, et al. 2020). Whereas effective TE suppression outside dynamic regions might be an important mechanism to restrict TE expansions in the core genome, the activity TEs in dynamic regions suggests that that these regions are prone to TE-mediated dynamics.

Environmental changes can alter suppression mechanisms and drive a de-repression of TEs (Bouvet, et al. 2008; Carr, et al. 2010; Fouche, et al. 2020). We observed TE expression upon changes in environmental conditions, i.e. different *in vitro* growth media as well as *in planta* growth. In addition, TE expression has been observed previously in different abiotic conditions in *V. dahliae* (Amyotte, et al. 2012; Faino, et al. 2016; Cook, et al. 2020). For example, some Tc1/Mariner and Mutator elements are induced upon heat stress (Amyotte, et al. 2012), while we observed that some LTR/Copia and LTR/Gypsy families highly expressed *in planta*. However, the function of TE de-repression under changing environmental conditions remains unknown (Galhardo, et al. 2007; Slotkin and Martienssen 2007; Koonin and Wolf 2009). Modifications in suppression patterns induced by changes in the environment could imply a trade-off between the functional TE de-repression and increased probability of TE expansions and TE mobilization on nearby genes, or even changes in the genome structure (Seidl, et al. 2016). Thus, although the role of environment and TE de-repression on genome function and structure remains unclear, the expression of dynamic TEs induced by changes in the environment could be an important driver of the genome dynamics in *V. dahliae*.

Changes in the TE landscape have often been associated with gene expression changes (Goubert, et al. 2020). Insertion of TEs within or near genes usually reduce gene expression, for example by spreading of silencing marks, such as DNA methylation or inducing changes in chromatin conformation (Castanera, et al. 2016; Winter, et al. 2018). TE insertions are associated with gene silencing and mostly have negative effects, yet also examples for an adaptive advantage exist. For example, in *Z. tritici* the insertion of a TE in the promoter of a melanin biosynthesis gene causes reduced melanin synthesis and quicker growth, which would be beneficial for colonization (Krishnan, et al. 2018). We observed that genes in proximity to TEs were typically less expressed during *in vitro* and *in planta* conditions, which was particularly clear for genes flanked by TEs. However, we observed that genes located in TE-rich regions are highly expressed *in planta*, raising a conundrum to link these opposing observations. Studies into dynamic genomic regions in *V. dahliae* and analogous regions in other fungi revealed that chromatin represents an additional layer of genomic regulation that creates a chromatin state allowing for accessibility to the transcriptional machinery (Schotanus, et al. 2015; Fokkens, et al. 2018; Soyer, et al. 2019; Cook, et al. 2020). We observed an enrichment of dynamic TEs in close proximity of these highly expressed genes in dynamic regions. This observation raises the possibility that dynamic TEs form part of the regulation of genes in the vicinity as *cis*-regulatory sequences or as a consequence of changes in the chromatin conformation (Le Rouzic, et al. 2007; Hollister and Gaut 2009; Huang, et al. 2012; Szitenberg, et al. 2016; Choi and Lee 2020). Furthermore, this could explain the maintenance of dynamic TEs in *V. dahliae*, as the TE suppression of these elements could negatively affect the expression levels of neighboring genes, favoring their co-localization.

Genome structure can differ significantly between species and even between strains of the same species (Lynch 2007). Here, we demonstrated that even closely related *V. dahliae* strains are characterized by their unique SV repertoire and co-localization of dynamic TEs and SVs in dynamic regions. As previously reported, we similarly observed high levels of sequence conservation in dynamic regions (de Jonge, et al. 2013; Faino, et al. 2016; Shi-Kunne, et al. 2018; Depotter, et al. 2019). Recently, dynamic regions were characterized by an intermediate chromatin state between heterochromatin and euchromatin (Cook, et al. 2020), which could indicate that this unique chromatin organization can allow for lower mutation rates in these dynamic regions (Prendergast, et al. 2007). We observed that SVs emerge rapidly due to their association with dynamic TEs and they represent a relevant mechanism to generate variation, rather than single-nucleotide changes in dynamic regions. The co-localization of TEs and SVs in distinct regions has been reported previously associated with genome compartmentalization in diverse fungi (Schotanus, et al. 2015; Faino, et al. 2016; Plissonneau, et al. 2016; Plissonneau, et al. 2018; Shi-Kunne, et al. 2018; Ola, et al. 2020; Torres, et al. 2020). Relaxed selection has been proposed to explain the association of variation and TEs in genomic compartments for example, the TE-rich accessory chromosomes in *Z. tritici* (Croll and McDonald 2012; Hartmann, et al. 2017; Grandaubert, et al. 2019). Conversely, dynamic regions in *V. dahliae* do not show signs of relaxed or strong positive selection (Depotter, et al. 2019). Since *V. dahliae* is an asexual fungus and consequently has a low effective population size, genetic drift could represent an important evolutionary force that maintains SVs in population (Kelkar and Ochman 2012; Cvijovic, et al. 2015), and these can therefore contribute to genome dynamics.

Compartmentalized genomes have been observed in diverse organisms (Torres, et al. 2020). In filamentous plant pathogens, this genome organization is described in the ‘two-speed’ genome model (Raffaele, et al. 2010) in which dynamic genome compartments serve as cradles of variation (Croll and McDonald 2012; Frantzeskakis, et al. 2019; Torres, et al. 2020). Additionally, changes in chromatin organization have been shown to be relevant for compartmentalized genomes to further increase or facilitate genomic variation (Schotanus, et al. 2015; Moller, et al. 2019; Cook, et al. 2020). Thus, further insights into chromatin organization and dynamic TEs are necessary to further disentangle the impact of chromatin modifications on the emergence of genetic variation and unravel how these facilitate the evolution of virulence in plant pathogens.

## Material and Methods

### *Verticillium dahliae* JR2 genome, repetitive element, and functional gene annotation

Repetitive element annotation of the chromosome-level genome assembly of *V. dahliae* strain JR2 (Faino, et al. 2015) was based on available annotation (Faino, et al. 2015; Cook, et al. 2020; Seidl, et al. 2020). Briefly, repetitive elements were annotated by using a combination of LTRharvest (Ellinghaus, et al. 2008), LTRdigest (Steinbiss, et al. 2009), and RepeatModeler. The repeat predictions were further curated and classified (Wicker, et al. 2007) by a combination of PASTEC (Hoede, et al. 2014) and sequence similarity to known transposable elements. The genome-wide occurrence of repeats was determined with RepeatMasker v 4.0.9, and the output was further processed using ‘one code to find them all’ (Bailly-Bechet, et al. 2014). Simple repeats and low-complexity regions were excluded (Cook, et al. 2020; Seidl, et al. 2020). GC-content, Kimura distance from a consensus TE family, and weighted DNA methylation data were previously summarized (Cook, et al. 2020; Seidl, et al. 2020). The composite RIP index (CRI) was calculated by obtaining the RIP substrate and the RIP product index as defined by nucleotide frequencies: RIP product index = TpA/ApT and the RIP substrate index= (CpA + TpG)/(ApC+GpT).

Functional gene annotation was based on the available *V. dahliae* JR2 gene annotation (Faino et al., 2015) and was performed using eggNOG 5.0 (Huerta-Cepas, et al. 2018) based on an hierarchical non-supervised orthology search, restricted to a fungal database with e-values >=0.001 and query coverage >=50%. To define the *V. dahliae* secretome, we predicted N-terminal signal peptides using SignalP v.4.1 (Nielsen 2017). Subsequently, we predicted putative effectors with EffectorP v.2.0 (Sperschneider, et al. 2018), using default parameters. Secondary metabolite biosynthetic genes were previously predicted using antiSMASH fungal version 4.0.2 (Weber, et al. 2015; Shi-Kunne, et al. 2018). Similarly, we used previously predicted CAZymes that cover signatures of glycoside hydrolases, polysaccharide lyases, carbohydrate esterases, glycosyltransferases, and carbohydrate-binding molecules (Seidl, et al. 2015; Shi-Kunne, et al. 2018). For further analysis, we considered the groups of ‘secreted proteins’ as the secreted proteomes excluding candidate effectors, secondary metabolite genes, and CAZymes.

### Gene and transposable element RNA-sequencing analysis

Transcriptional activity of genes and repetitive elements in *V. dahliae* strain JR2 was assessed using previously generated transcriptome data (Cook, et al. 2020) of three *in vitro* conditions, (Potato Dextrose Broth, Murashige-Skoog and Czapeck-Dox media) and of *in planta* colonization (*Arabidopsis thaliana*; 28 dpi). For further analysis, we considered only MS media for *in vitro* and *in planta* comparisons. Single-end sequencing reads of three biological replicates per condition were mapped to *V. dahliae* strain JR2 genome assembly (Faino, et al. 2015) using STAR v.2.4.2.a, allowing multiple mapped reads with the following settings: ╌outFilterMultimapNmax 100 ╌ winAnchorMultimapNmax 200 ╌outSAMtype BAM Unsorted ╌outFilterMismatchNmax 3 (Jin, et al. 2015; Jin and Hammell 2018; Fouche, et al. 2020). The bam files were sorted by read name with samtools v.1.2 and the transcriptional activity level was quantified using TEcount from the TEtrancripts package 2.2.1 (Jin, et al. 2015), with the following parameters: ╌stranded no –mode multi, -iteration 1000. TEcount considers multiple-mapped reads aligned to genes and TE regions to determine transcript abundance per condition/replicate. Furthermore, TEcount only considers reads that map in its entirety to TEs and reads mapping to only a fraction were discarded (Jin, et al. 2015; Jin and Hammell 2018).

Sequencing reads summarized over TEs and genes were normalized between three replicates in the different conditions using R/Bioconductor package EdgeR v.3.8 (Robinson, et al. 2010; Robinson and Oshlack 2010). For normalization, we only considered genes and TEs >=1 reads in all samples (Anders, et al. 2013). TEs with read count 0 were assumed to have no transcriptional activity. Libraries were normalized with TMM method (Robinson and Oshlack 2010), and converted to counts per million (CPM) mapped reads using R/Bioconductor package EdgeR v.3.8 (Robinson, et al. 2010). Comparisons between transcriptional levels were computed in R 3.6.3 (Team 2019)

### Single nucleotide variant detection and analysis

Single nucleotide variants were detected using paired-end sequencing reads of each 42 previously sequenced *V. dahliae* strains (Supplementary Table 1). Each strain was mapped independently to the reference genome *V. dahliae* JR2 using BWA -MEM v.0.7 with default options (Li and Durbin 2010). Library artifacts were marked and removed using Picard tools v.2.18 with -MarkDuplicates followed by -SortSam to sort the reads (http://broadinstitute.github.io/picard/). Single nucleotide variants were identified using the -HaplotypeCaller of the Genome Analysis Toolkit (GATK) v.4.0 (Poplin, et al. 2018). Variations were detected individually per strain, using -ploidy 1 and -emitRefConfidence GVCF. Then, all strains were joined by -Jointgenotyping with -maxAltAlleles 2, and all non-SNVs were removed using SelectVariants -selectType SNP. To obtain high-quality SNVs, we applied the following cut-off filters: QUAL<250, MQ<40, QD<20, FS>60, SOR>3.0, ReadPosRankSum<-5.0, MQRankSum between −2 and 2. Finally, we excluded variants with missing genotype calls in 10% of the strains. To explore the similarity between strains, an unrooted phylogenetic tree was constructed using a Neighbor-Joining approach based on the final single nucleotide variance set, using distance function in R 3.6.3 (Team 2019).

To further assess the sequence diversity between *V. dahliae* strains, we calculated the nucleotide diversity (π) based on Nei and Jin (1989) using a sliding window of 1 kb (500 bp sliding) as implemented in the PopGenome package (Pfeifer, et al. 2014) in R. Single nucleotide variants were annotated using SNPeff v.3.2 (Cingolani, et al. 2012) using the refined annotation of *V. dahliae* JR2 strain. Based on the SNPeff prediction, we calculated the number of synonymous and non-synonymous mutations.

### Structural variant calling and analysis

To predict structural variants, we used the ‘sv-callers’ workflow with few modifications that enabled parallel execution of multiple SV callers (Kuzniar, et al. 2020). We used four different callers: DELLY v.0.8.1 (Rausch, et al. 2012), LUMPY v.2.13 (Layer, et al. 2014), Manta v.1.6.0 (Chen, et al. 2016), and Wham v.1.0 (Kronenberg, et al. 2015) in single-sample mode, using each 42 previously sequenced *V. dahliae* strains independently (Supplementary Table 1). Briefly, the workflow takes every strain and maps the genomic reads to the reference genome *V. dahliae* JR2 using BWA-MEM v.0.7 with default options (Li and Durbin 2010). Then, library artifacts were marked and removed using Picard tools v.2.18 with -MarkDuplicates followed by -SortSam to sort the reads (http://broadinstitute.github.io/picard/). The four SV callers were used with default parameters, except for DELLY in which we used >1 as minimum quality for further processing. All outputs were first filtered using bcftools -filter v.1.3.2 with default settings used in the ‘sv-callers’ workflow (Li 2011), except for Manta since the score model for post-processing assumes a diploid genome (Chen, et al. 2016). Therefore, we used the unscored Manta predictions for further processing.

For final filtering, the ‘sv-callers’ workflow post-processes the vcfs. Briefly, the results of the four different callers per strain were merged using SURVIVOR v.1.0.6 (Jeffares, et al. 2017), keeping SVs predicted by at least three callers, allowing 1000 bp as the maximum distance from breakpoints predicted by the different tools and considering only same SV types. Subsequently, only SVs with minimum size >50 bp and maximum size 1 Mb as well as localization outside of a low-quality region, defined as MQ=0 and read support <10 for every strain, were kept. Finally, the 42 independent datasets we combined into a single vcf file using SURVIVOR by merging SVs up to 1000 bp apart of the same SV type.

### Transposable element polymorphism prediction and analysis

Transposable element presence/absence polymorphisms were analyzed using TEPID v.2.0 (Stuart, et al. 2016), using paired-end short reads of 42 previously sequenced *V. dahliae* strains (Supplementary Table 1). TEPID integrates split and discordant read mapping, mapping quality, sequencing breakpoints, and local variations in coverage to identify variants with respect to a reference TE annotation (Stuart, et al. 2016). We mapped each of the 42 *V. dahliae* strains individually to *V. dahliae* JR2 reference genome, using tepid-map (average insert size −s 500), which uses Bowtie2 v.2.2.5 (Langmead and Salzberg 2012). TE variant discovery was computed using tepid-discover, considering insertion and deletion prediction and a conservative discovery through –strict option. We subsequently refined the discovered variants using tepid-refine to reduce false-negative calls within the group of 42 strains.

### Statistical analysis and visualization

The distribution of SVs over the *V. dahliae* strain JR2 genome was determined considering the breakpoints (±1 bp) overlapping in 10 kb non-overlapping windows. We predicted an expected probability distribution assuming a Poisson distribution, using λ=0.77 (mean number of SVs breakpoints per 10 kb windows). To investigate if SVs co-localize, we performed a clustering analysis using CROC (Pignatelli, et al. 2009) based on a hypergeometric distribution test and posterior multiple-testing correction using Benjamini-Hochberg; CROC scans every chromosome with a sliding window (SV-breakpoints per window=10, slide=1, >3 SV-breakpoints as a minimum). We computed the same test for TE clustering using TEs per window=10, slide=1, >3 dynamic TEs as a minimum. Permutation tests were computed using R/Bioconductor regionR v.1.18.1 package (Gel, et al. 2016) and performed 10,000 iterations, using the mean distance to evaluate the closest relationship (bp distance) between SV-breakpoints and TE elements, and circular randomization to maintain the order and distance of the regions in the chromosomes. We assumed the same parameters to evaluate the association of TEs and LS regions but considering the number of overlaps instead of distance. Finally, we computed enrichment association tests using bedtools -fisher v.2.25.0 based on a one-sided Fisher’s test (Quinlan and Hall 2010).

Principle component analyses were performed in R v.3.6.3, using packages FactoMineR v.1.42 and factoextra v.1.0.5 (Lê, et al. 2008). The used variables for each TE were methylation (mCG, mCHG, mCHH), GC content, Kimura distance, CRI, Frequency, and *in vitro* (MS media) expression. To further investigate the relationship of TEs with gene expression, we summarized TEs in 5kb windows upstream and downstream genes, using bedtools -window (Quinlan and Hall 2010). Then, we classified genes in upstream, downstream if >=1 TE in each independent position. We considered genes in between TEs as ones with >=1 TEs in both positions and we exclude them from the other categories. We ranked each gene near TE based on their expression profile and counted the number of TEs associated with each independent rank. These rank counts were then binned to obtain the distribution of ranks and fit a linear model for the data and calculated the R^2^ and P value for the fit of the model in R v.3.6.3 (Team 2019). All statistical analyses and comparison tests were performed in R v.3.6.3 (Team 2019), and visualization with R packages ggplot2 and circlize (Gu, et al. 2014; Wickham 2016).

## Supporting information

SFig1

SFig2

SFig3

SFig4

SFig5

SFig6

SFig7

SFig8

SFig9

SFig10

SFig11

STabkes1-10

## Acknowledgments

This work was supported by the Consejo Nacional de Ciencia y Tecnología, México and the Deutsche Forschungsgemeinschaft (DFG, German Research Foundation) under Germany’s Excellence Strategy – EXC 2048/1 – Project ID: 390686111.

## Data availability statement

The data underlying this article are accessible at the National Center for Biotechnology Information (NCBI) Sequence Read Archive (SRA) under BioProject PRJNA592220, and mentioned in Supplementary Table 1.

