## Supplementary legends for "Transposable elements contribute to genome dynamics and gene expression variation in the fungal plant pathogen *Verticillium dahliae*"

**Supplementary Figure Legends**

**Supplementary Figure 1. Structural variant (SV) calling methods used in this study. (A)** The total number of SV predictions from the four different SV callers and their overlap is shown as a Venn diagram; **(B)** Overview of the 3,530 SVs identified in the 42 *V. dahliae* isolates after combining the results with SURVIVOR. For the ‘final set’, we only considered SVs predicted by at least three different callers (see Methods) and removed SVs due to mapping in low-quality regions (MQ=0) and to size (>1 Mb) (see Methods). The ‘final set’ was considered for all further analyses; **(C)** Length distribution of the final set of 505 SVs, here excluding translocations, used for all further analysis.

**Supplementary Figure 2. Distribution of SV breakpoints in the *Verticillium dahliae* genome.** Density plot overlapping the empirical and expected probability distribution in 10 kb non-overlapping windows across the genome of *V. dahliae* strain JR2. The empirical distribution considers the breakpoints of all 941 SVs. The expected probability distribution was estimated assuming a Poisson distribution. Statistical difference between the two distributions was assessed using a Kolmogorov-Smirnov test (P<2.2e-16).

**Supplementary Figure 3.** **Dynamic genomic regions in *Verticillium* *dahliae* are enriched in structural variants (SVs). (A)** Enrichment of different classes of SVs in core and dynamic genomic regions. The heatmap displays the posterior false-discovery rate corrected p-value derived from a Fisher´s exact test; **(B)** Length (log10) distributions of SVs overlapping the core and dynamic genomic regions. The statistical difference has been assessed using a one-sided Wilcoxon rank-sum test.

**Supplementary Figure 4.** **Transposable elements are close to SVs. (A)** Permutation test to assess the distance between the full set of TEs and SV breakpoints. The vertical dashed line represents the mean distance of overlaps expected at 10,000 random permutations, the yellow line indicates the mean distance observed at a closer distance (11,284 bp), significance P = 9.9e-5, z-score=-5.43. TEs showed to be more closed to SV breakpoints than expected by chance; **(B)** The distribution of TE density (TE counts) summarized in 10 kb non-overlapping windows is shown. TEs at centromeric regions were removed. The statistical significance was assessed using a Wilcoxon rank sum test.

**Supplementary Figure 5.** **Principle component analyses of TEs in *V. dahliae*.** Decomposition of the principal component analysis for eight variables summarized for each TE, excluding centromeric regions. Each vector represents one variable, with the length indicating the importance of the variable for this dimension, for angles <90º, the two variables are correlated, while >90º indicates that the variables are negatively correlated. The coordinates and contribution for each variable are further detailed in Supplementary Table 4.

**Supplementary Figure 6. Transcript levels of *V. dahliae* strain JR2 TEs in two *in vitro* conditions.** TE expression is displayed for two *in vitro* conditions, PD (Potato Dextrose broth) and CZ (Czapek-Dox) growth media. Expression values are log2(CPM+1) transformed. p-value after one-sided Wilcoxon rank-sum test.

**Supplementary Figure 7. TE insertions are not enriched in dynamic genomic** **regions in *V. dahliae* strain JR2. (A)** Permutation test of overlaps between new TE insertions and dynamic genomic regions. The vertical dashed line represents the mean number of overlaps expected at 10,000 random permutations and the red line indicates the number of overlaps observed (7), significance P= 0.451, z-score=-0.534; **(B)** Permutation test of overlaps between original coordinates in JR2 of TE insertions and dynamic genomic regions. The vertical dashed line highlights the mean number of overlaps expected at 10,000 random permutations and the red line indicates the number of overlaps observed (18), significance P= 0.294, z-score=0.7862.

**Supplementary Figure 8. TE distribution relative to protein-coding genes in *V. dahliae*.** Density distribution of non-dynamic TEs depicted by superfamilies relative to the location of protein-coding genes (distance=0), showing a random distribution within 5 kb windows upstream and downstream gene sequences.

**Supplementary Figure 9. TE expression under different conditions.** Unsupervised clustering of TE expression monitored under different *in vitro* conditions (Czapek-Dox, CZ; potato dextrose broth, PD; Murashige-Skoog, MS) and *in planta* condition (*Arabidopsis* 28 dpi; AR). The mean transcript abundance (counts per million; log2(CPM+1) transformed) is depicted in the heatmap. The blue-coded left column details the superfamily, and the yellow-coded column summarizes the abundance of dynamic TEs in each of the TE family.

**Supplementary Figure 10. Gene expression correlates with transposable elements occurrence (A)** *In planta* (Arabidopsis 28 dpi) gene expression of genes located upstream (*n*=1,098), downstream (*n*=1,051), between (*n*=546) TEs (5 kb proximity), or genes without TEs in proximity = Not close (5 kb; *n*=8,742) are shown; **(B)** *In vitro* and (**C**) *in planta* gene expression of genes upstream (*n*=153), downstream (*n*=134), and between (*n*=118) dynamic TEs; **(D)** Comparison of distances between dynamic TEs to genes (within 5 kb windows); in all cases, p-value depicts one-sided Wilcoxon rank-sum test.

**Supplementary Figure 11. Dynamic transposable elements correlate with gene expression depending on their position on the genome.** Relationship between gene expression ranks (percentile windows, from low (blue) to high (red) expression; *in vitro* and *in planta*) and TE density, depending on relative TE positions (within 5 kb). The linear regression (dark line) and confidence interval (light grey) are shown as well as the R and p-value after linear regression. The upper panels depict the dynamic TEs and the lower panels the non-dynamic TEs.
