## Supplementary figures and images for "Transposable elements contribute to genome dynamics and gene expression variation in the fungal plant pathogen *Verticillium dahliae*"

### SFig1

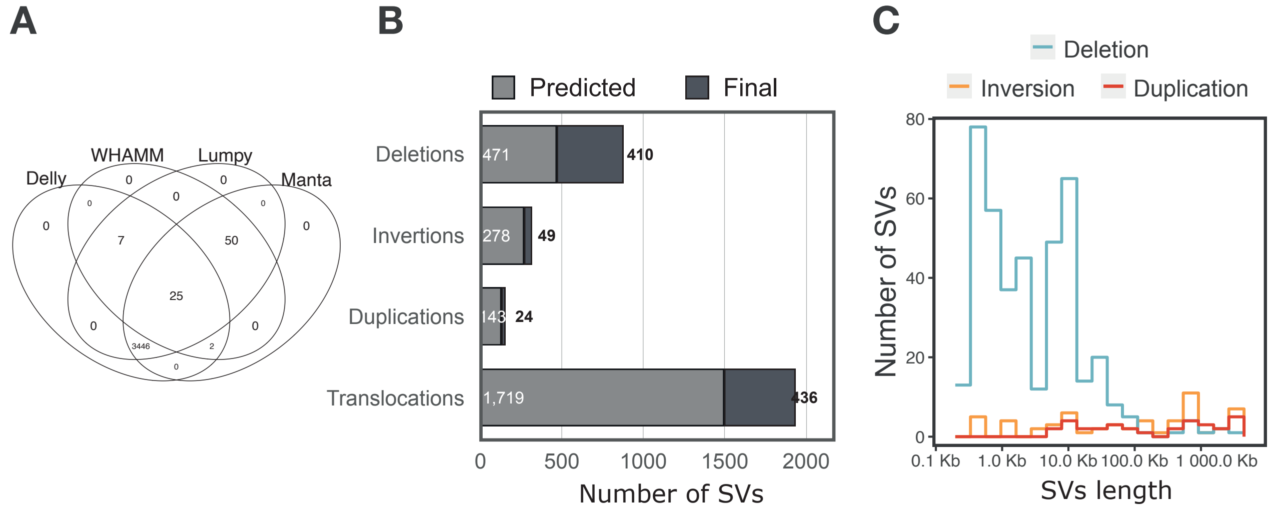

### SFig2

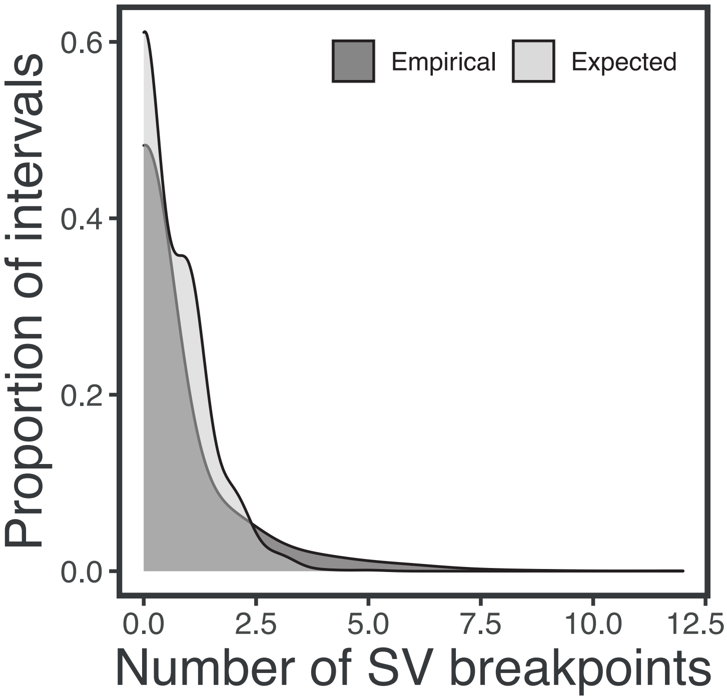

### SFig3

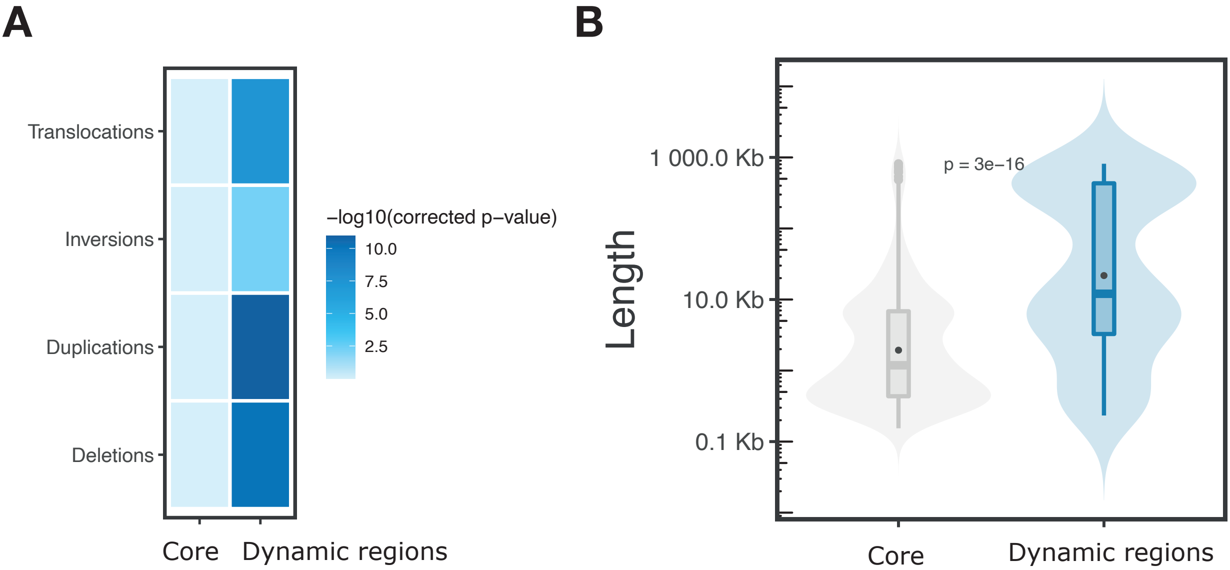

### SFig4

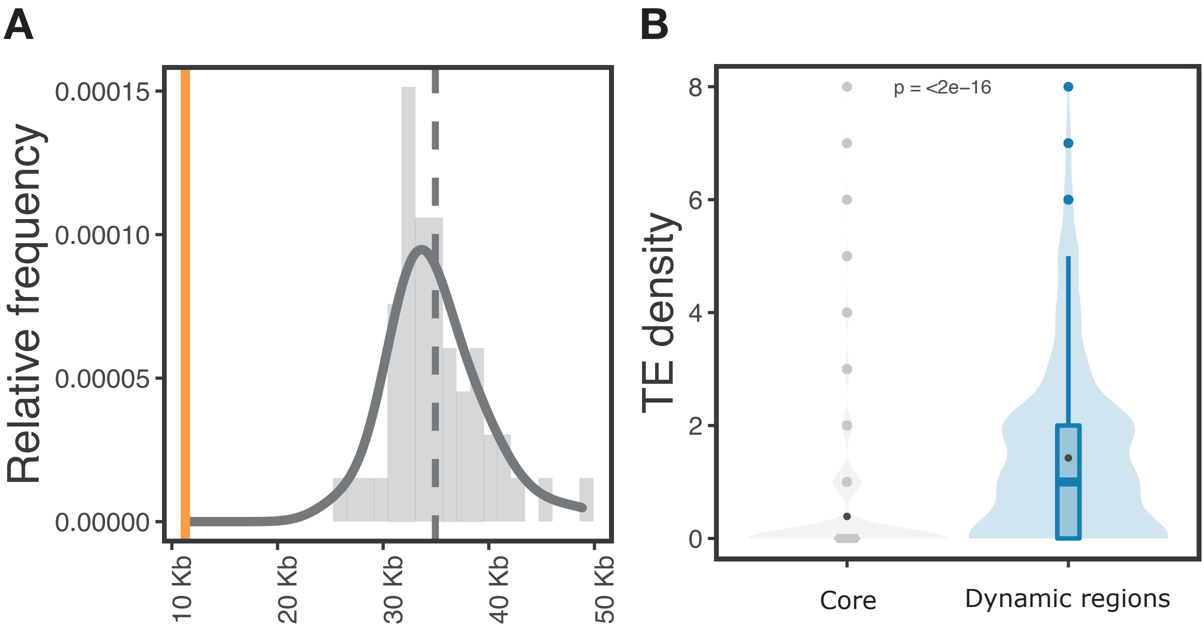

### SFig5

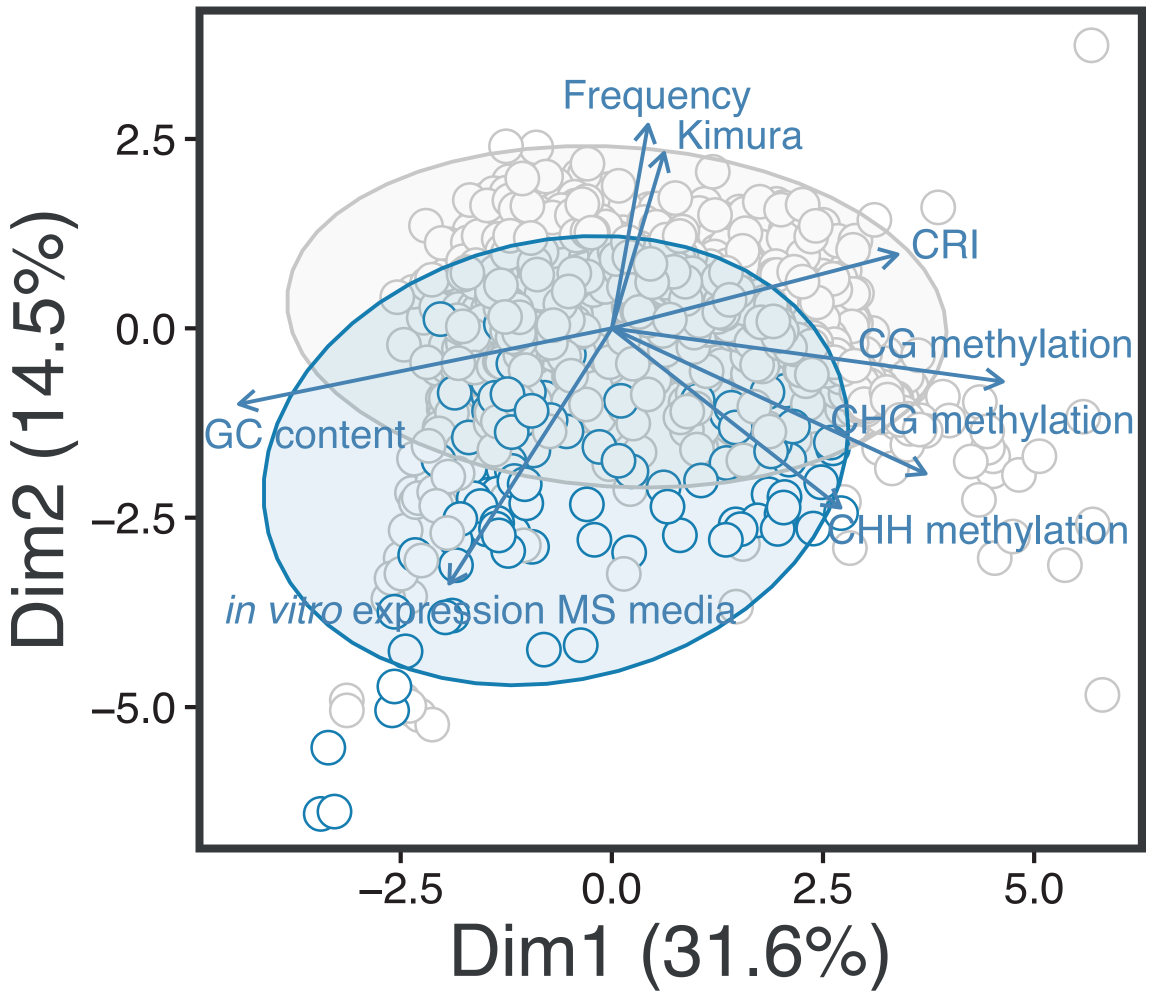

### SFig6

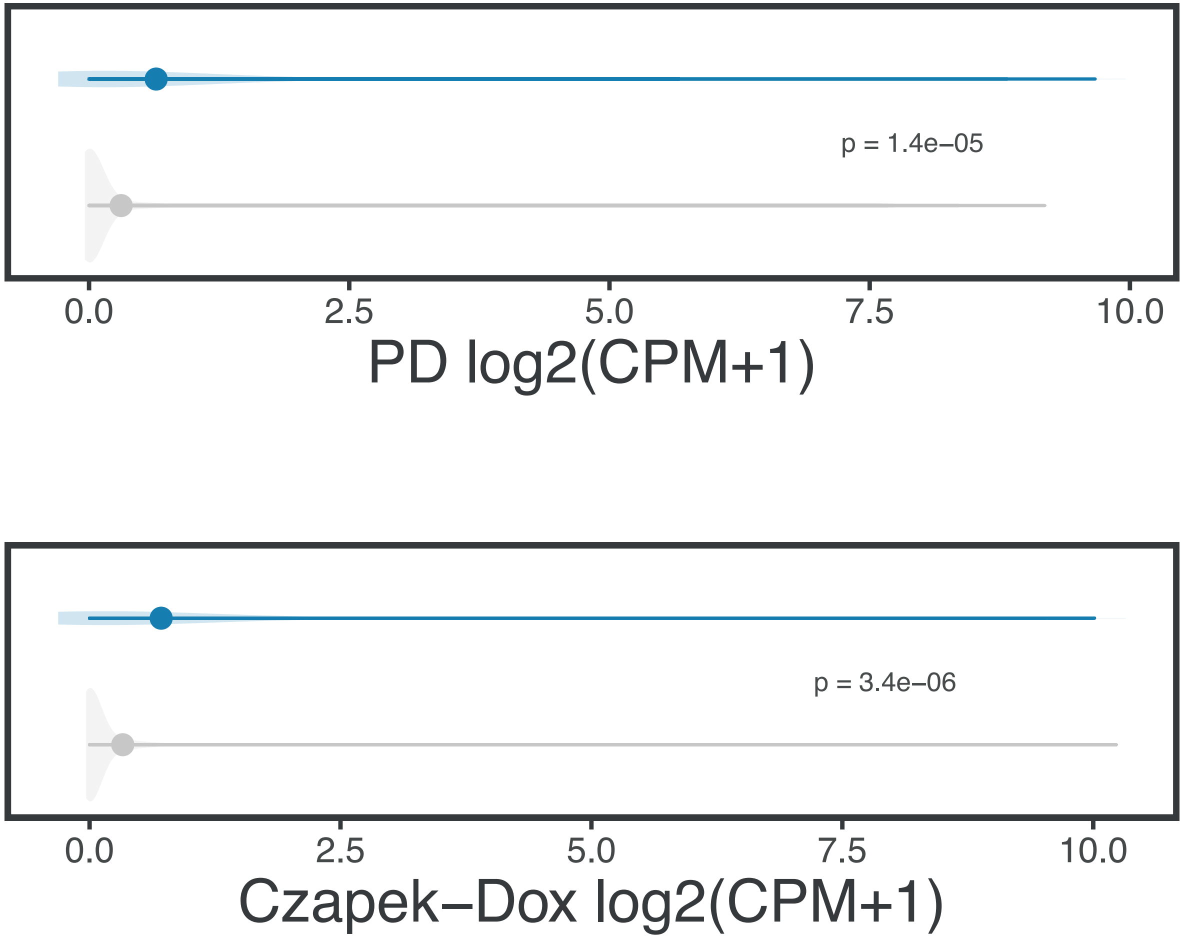

### SFig7

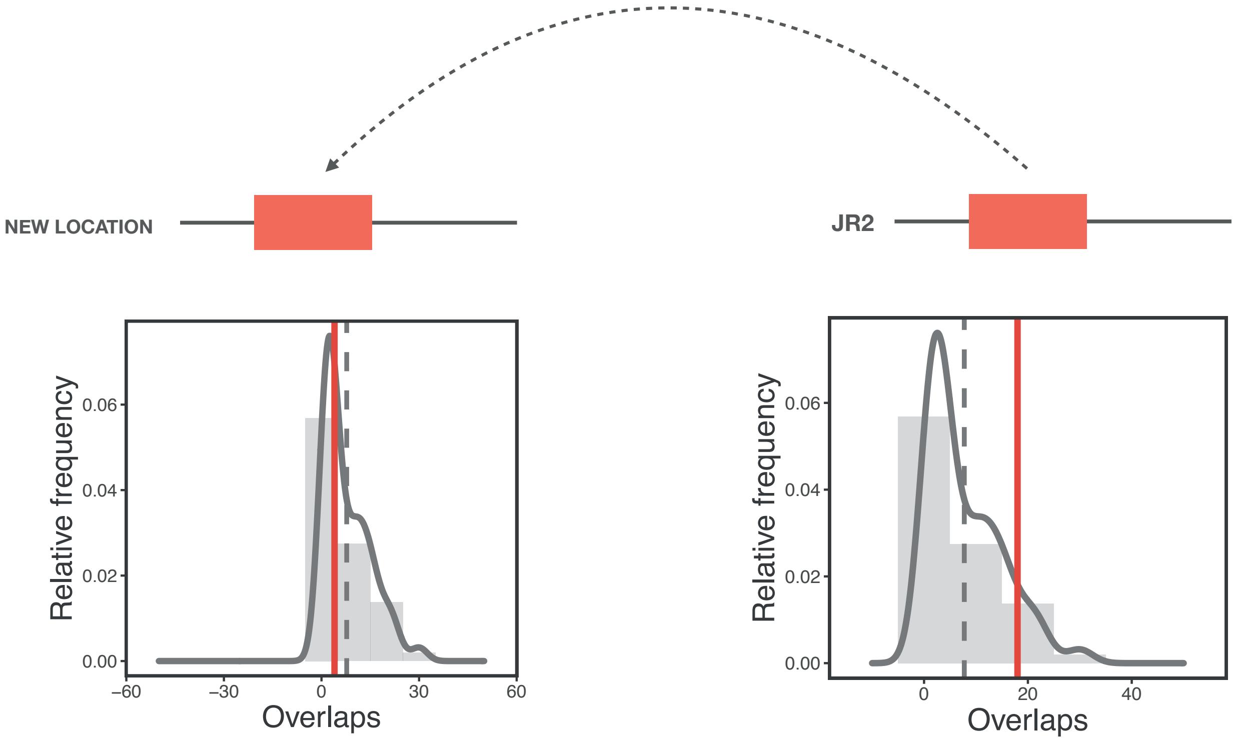

### SFig8

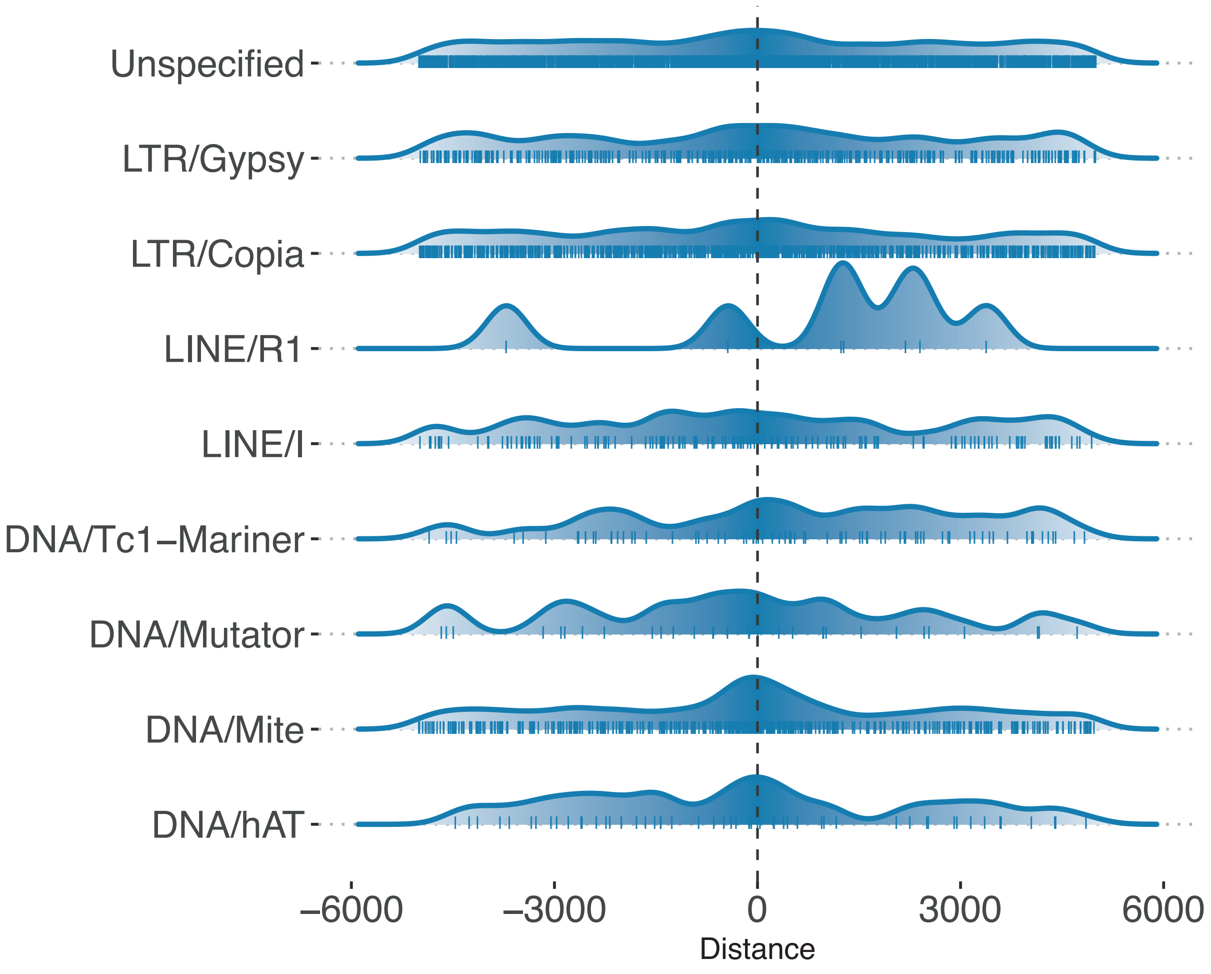

### SFig9

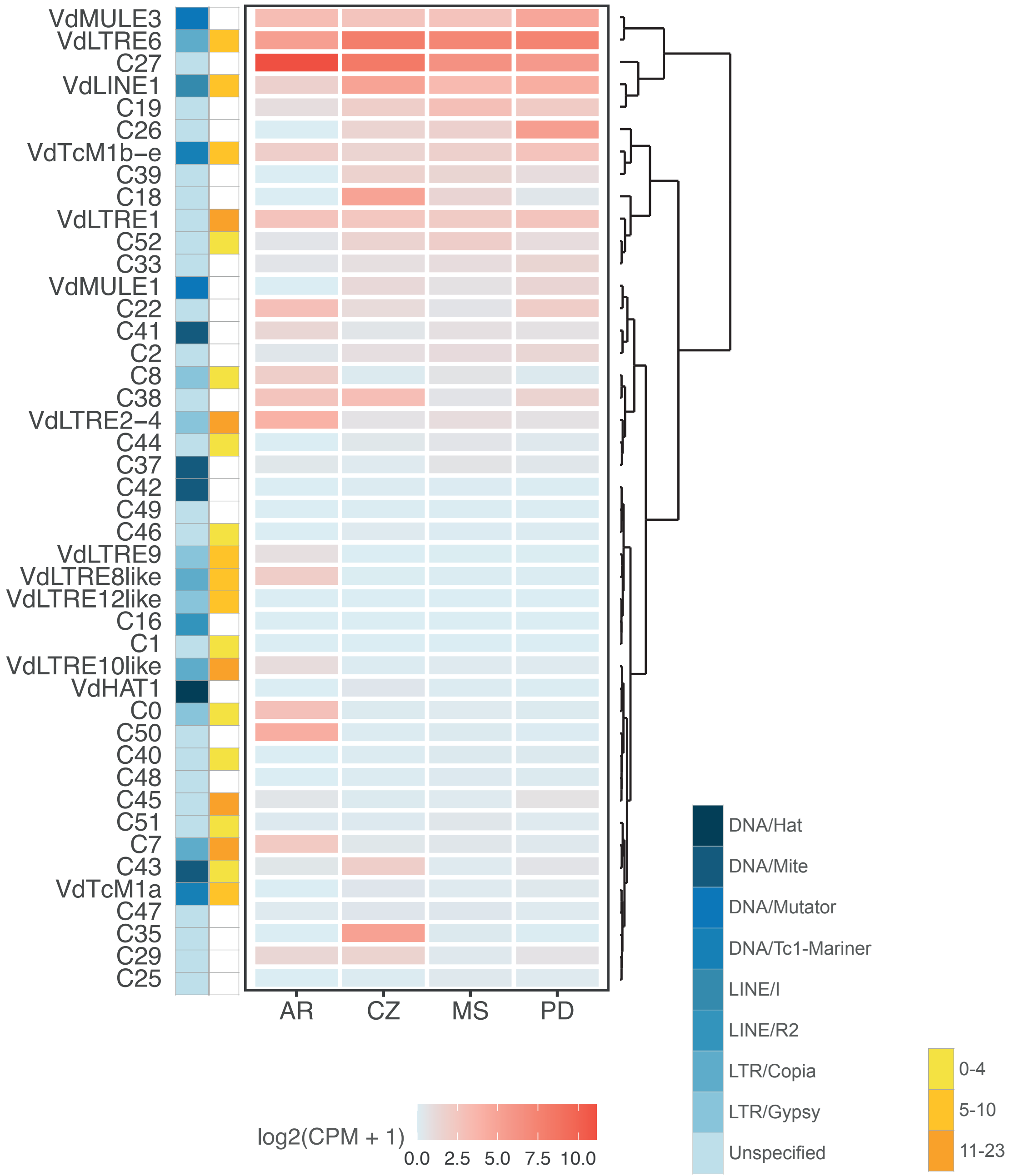

### SFig10

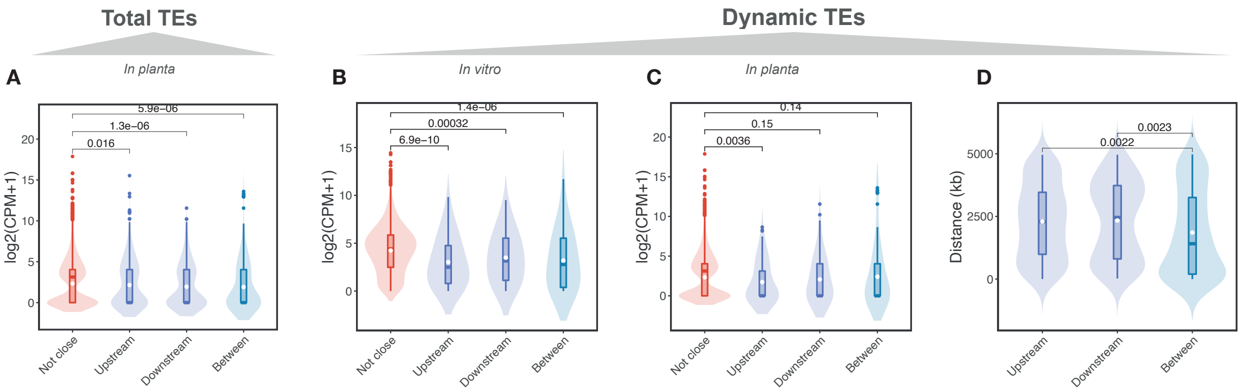

### SFig11

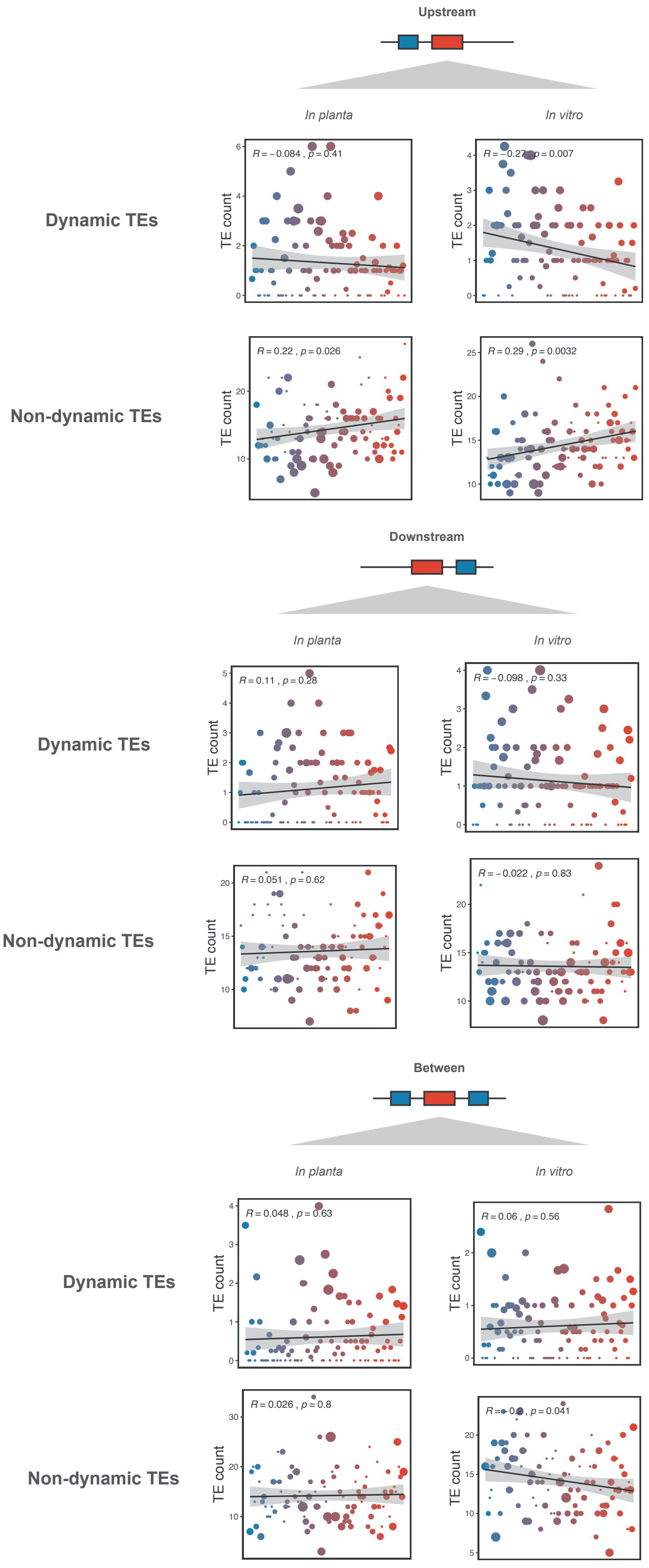
